# Putative neofunctionalization of a Poales-specific EXO70 clade

**DOI:** 10.1101/2024.12.13.628418

**Authors:** Molly Bergum, Jan Sklenar, Inmaculada Hernández-Pinzón, Jodie Taylor, Matthew Smoker, Sebastian Samwald, Megan Allen, Anupriya Thind, Phon Green, Hyeran Moon, Frank L. H. Menke, Cyril Zipfel, Jack Rhodes, Christine Faulkner, Matthew J. Moscou

## Abstract

EXO70s are uniquely expanded in land plants compared to all other eukaryotic lineages. The functional implications of this expansion and diversification on the conserved role of EXO70 as a subunit of the octameric exocyst complex have remained unresolved. We previously demonstrated barley (*Hordeum vulgare*) EXO70FX12, a member of the monocot-specific EXO70FX clade, is required for resistance to wheat stripe rust in conjunction with the leucine-rich repeat receptor kinase (LRR-RK) HvPUR1. Through phylogenetic analysis, we identified unique features of the EXO70FX clade, leading us to hypothesize that this clade experienced neofunctionalization. Using structural predictions and protein-protein interaction assays, we demonstrate that HvEXO70FX12 lost the ability to serve as a subunit within the exocyst complex. We predict that the EXO70FX clade has largely lost exocyst association and represents a novel acquisition that emerged during Poales diversification for immunity.

## INTRODUCTION

The octameric exocyst, consisting of SEC3, SEC5, SEC6, SEC8, SEC10, SEC15, EXO84, and EXO70, is an evolutionarily conserved complex found across eukaryotic lineages (TerBush and Novick 1995; TerBush et al. 1996; Dong et al. 2005; Cvrčková et al. 2012; Boehm and Field 2019). The exocyst is involved in trafficking secretory vesicles in a variety of critical processes in yeast, mammals, and plants, including polarised exocytosis and cytokinesis (Wu and Guo 2015). During targeted secretion, SEC3 and EXO70 bind to phospholipids to anchor the complex to the plasma membrane (PM) while other subunits, including SEC15 and SEC6, interact with the trafficked vesicles prior to SNARE-mediated fusion (Guo et al. 1999; He et al. 2007; Zhang et al. 2008; Shen et al. 2013; Wu and Guo 2015). While SEC3 was shown to have a dominant role in localising the exocyst in yeast and mammals, EXO70A1 was shown to primarily mediate this localised targeting in *Arabidopsis thaliana* (Synek et al. 2021).

EXO70, first identified in yeast as a component of the exocyst complex (TerBush et al. 1996), is uniquely expanded in plants (Cvrčková et al. 2012; Boehm and Field 2019). Whilst existing as a single copy in other eukaryotes, EXO70s in green plants have evolved in three monophyletic families, EXO70.1, EXO70.2, and EXO70.3, comprising eight conserved clades in angiosperms, designated with a letter suffix from A to H (Synek et al. 2006, 200; Žárský et al. 2020). EXO70s from diverse clades function in plant immunity across monocots and dicots. AtEXO70B1 (Stegmann et al. 2013; Wang et al. 2019, 2020), AtEXO70B2 (Pečenková et al. 2011; Stegmann et al. 2012), AtEXO70H4 (Huebbers et al. 2024), OsEXO70B1 (Hou et al. 2020), OsEXO70E1 (Guo et al. 2018), OsEXO70H3 (Wu et al. 2022), OsEXO70F3 (Fujisaki et al. 2015), HvEXO70FX11b (Ostertag et al. 2013), and HvEXO70FX12 (Holden et al. 2022) have each been implicated in regulating resistance to pathogens or insects in *A. thaliana*, rice (*Oryza sativa*), or barley (*Hordeum vulgare*).

The plant immune system is comprised of pathogen recognition that occurs both inside and outside the plant cell (Jones and Dangl 2006). Spanning the PM, pattern-recognition receptors (PRRs), which include receptor kinases (RKs) and receptor proteins (RPs), perceive extracellular microbial or modified-self patterns to induce pattern-triggered immunity (PTI) (Couto and Zipfel 2016). Inside of the cell, immune signalling is largely elicited by nucleotide-binding domain leucine-rich repeat (NLR) proteins, which respond to microbial effectors and cause effector-triggered immunity (ETI) (Jones and Dangl 2006; Adachi et al. 2019). While receptor activation differs between PTI and ETI, both responses are mutually potentiated and can lead to immune responses, including apoplastic ROS bursts, cytosolic Ca^2+^ influx, MAPK signalling, transcriptional reprogramming, hormonal signalling, cell death, and secretion of defence compounds for cell wall reinforcements and pathogen antagonism (Bigeard and Hirt 2018; Ngou et al. 2021; Yuan et al. 2021a, 2021b; Bender and Zipfel 2023; Jian et al. 2023).

The mechanism of most EXO70s involved in plant immunity is predicted to require the exocyst, as EXO70s from A, B, E, and H clades have been shown to interact with exocyst subunits, and the immune mechanisms often include secretion of components to the PM or exocytosis (Pečenková et al. 2011; Kulich et al. 2013; Ding et al. 2014; Synek et al. 2021; Michalopoulou et al. 2022; Wu et al. 2022). AtEXO70B1 and AtEXO70B2 interact with both the exocyst and the PRR FLAGELLIN-SENSING 2 (AtFLS2), are required for AtFLS2 signalling, and promote the accumulation of multiple RKs including AtFLS2, BR INSENSITIVE 1 (AtBRI1), and CHITIN ELICITOR RECEPTOR KINASE 1 (AtCERK1), at the PM (Pečenková et al. 2011; Kulich et al. 2013; Wang et al. 2020). Additionally, AtEXO70B1 is the target of diverse effectors from bacterial pathogens, including XopP from *Xanthamonas campestris*, AvrPtoB from *Pseudomonas syringae* pv. tomato DC3000, and RipE1 from *Ralstonia solanacearum* (Wang et al. 2019; Michalopoulou et al. 2022; Tsakiri et al. 2022; Kotsaridis et al. 2023). Interestingly, XopP mediates virulence by preventing the association of AtEXO70B1 in the exocyst and subsequently impairing the exocytosis of PATHOGENESIS-RELATED GENE 1A (PR1A), deposition of callose, and localisation of AtFLS2 to the PM (Michalopoulou et al. 2022).

AtEXO70H4 is required for the polar secretion of callose synthases in the trichome, and AtMLO6, a member of a protein family that includes calcium channels, has a predicted role in targeting AtEXO70H4 to the PM (Kulich et al. 2018; Gao et al. 2023; Huebbers et al. 2024). Likely through a mechanism of vesicle trafficking, AtEXO70H4 and AtMLO6 confer susceptibility against powdery mildew (Huebbers et al. 2024). OsEXO70E1 and OsEXO70H3 are also expected to have an exocyst-dependent mechanism for broad-spectrum resistance against brown and white-backed planthoppers in rice mediated by a leucine-rich repeat (LRR) domain-containing protein, BPH6 (Guo et al. 2018; Wu et al. 2022). OsEXO70H3 interacts with various exocyst subunits and is required for reinforcing the cell wall via lignin deposition mediated by its interaction with the protein SAMSL (Wu et al. 2022).

It is unsurprising that other exocyst subunits have a similar importance in immunity, which can be attributed to the role of the exocyst in regulating the secretion of defence compounds or the deposition of callose (Du et al. 2018). In *Nicotiana benthamiana*, SEC5, SEC6, and SEC10 positively regulate resistance to the hemi-biotrophic pathogens *Phytophthora infestans* and *P. syringae* and, conversely, susceptibility to the necrotrophic pathogen *Botrytis cinerea* (Du et al. 2018). Furthermore, an RXLR effector from *P. infestans* interacts with potato (*Solanum tuberosum*) SEC5 to suppress callose deposition and enhance susceptibility (Du et al. 2015). More broadly, many components of the vesicle trafficking pathway are targeted by *P. infestans* PexRD12/31 effectors (Petre et al. 2021).

To date, it has been unknown whether an EXO70 involved in plant immunity could be acting independently of the exocyst complex. We have previously shown that *HvExo70FX12* is required for wheat stripe rust (*Puccinia striiformis* f. sp. *tritici*) resistance in barley (Holden et al. 2022). Immunity is only conferred in the presence of both *HvExo70FX12* and the gene encoding the sub-family XII leucine-rich repeat RK (LRR-RK) PUCCINIA STRIIFORMIS RK 1 *HvPur1*, which is located approximately 160 kb distal from *HvExo70FX12* in the barley genome (Holden et al. 2022). As the EXO70FX clade is only found in Poales species, relatively little is known about their function. Although the mechanistic connection between HvPUR1 and HvEXO70FX12 remains unsolved, evidence suggests that the role of HvEXO70FX12 in immunity is independent from the exocyst. Using phylogenetics, structural predictions, and protein-protein interaction assays, we predict the Poales-specific EXO70FX clade has undergone neofunctionalization.

## RESULTS

### EXO70s are highly diverse in green plants with lineage-specific expansions

Using recently released genomes of 23 green plant species, we constructed a maximum likelihood phylogenetic tree of EXO70 proteins utilising a structure-guided multiple sequence alignment with the outgroup species *Mus musculus* (mouse) and *Saccharomyces cerevisiae* (yeast). The basal lineage includes single EXO70s from *Mesostigma viride* and *Chlorokybus atmophyticus* of the earliest diverging Streptophyta lineage *Charophyta* (green algae) (Leliaert et al. 2011). The phylogenetic tree largely supports previous work that established three major lineages of EXO70: EXO70.1, EXO70.2, and EXO70.3 (Fig. 1; Synek et al. 2006; Cvrčková et al. 2012). While the EXO70.1 lineage lacks bootstrap support, all species were present including the single EXO70 from *Klebsormidium nitens* and two EXO70s from *Chara braunii*. Four EXO70s from *Mesotaenium endlicherianum* are found basal to all three major lineages, whereas the six EXO70s from *Spirogloea muscicola* are found basal to EXO70.1 and EXO70.3. Previous work on the evolution of the angiosperm EXO70 gene family established eight conserved EXO70 clades (A, B, C, D, E, F, G, H), the arbuscular mycorrhizal symbiosis associated EXO70I clade (Zhang et al. 2015), the lineage-specific EXO70FX clade in the Poaceae (grass) family (Synek et al. 2006; Holden et al. 2022), and the lineage-specific EXO70J clade in the Fabaceae (legume) family (Chi et al. 2015). Phylogenetic congruence was observed for the majority of clades including basal gymnosperm species with bootstrap support for EXO70C, EXO70D, EXO70E/EXO70F, EXO70G, and EXO70H. Poor bootstrap support was observed in the EXO70A clade, although this may be due to the inclusion of highly divergent basal lineages, and EXO70B lacks a clear gymnosperm ortholog. Large lineage-specific expansions in all three EXO70 lineages were observed in *Ceratopteris richardii* (fern), whereas a single expansion was observed basal to the EXO70H clade in *Picea abies* (Norway spruce) and *Pinus taeda* (Loblolly pine).

**Fig. 1.**
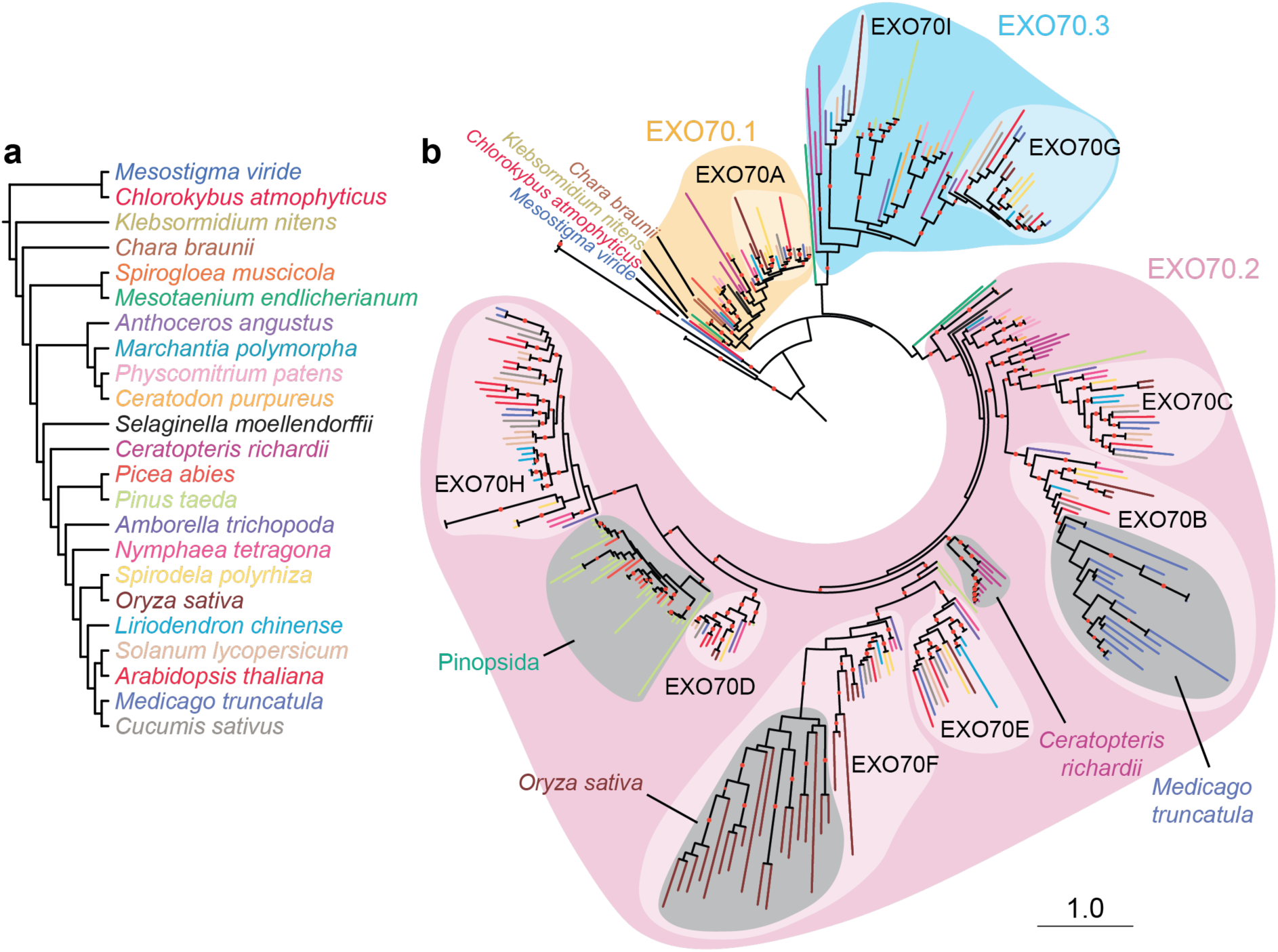
Green plant EXO70s are highly diverse and experience lineage-specific expansions. a) Phylogenetic relationship of 23 representative green plant species with colour-coding corresponding to branches in panel b (Leliaert et al. 2011). b) Maximum likelihood phylogenetic tree of 327 green plant EXO70s based on a structure-based alignment with EXO70s from yeast (PDB 2B1E and 5YFP), mouse (PDB 2PFT), and *A. thaliana* (PDB 4RL5). Previously described major lineages EXO70.1, EXO70.2, and EXO70.3 and angiosperm-specific clades A, B, C, D, E, F, G, H, and I are highlighted (Synek et al. 2006; Cvrčková et al. 2012). Lineage-specific expansions are observed in *Oryza sativa* (Poales; EXO70FX) and *Medicago truncatula* (Fabaceae; EXO70BX). Bootstrap support of greater than or equal to 80% is shown at branch midpoints with a red dot. Scale indicates 1.0 substitution per site.

Recent reclassification in the literature has created conflicting clade nomenclature for the EXO70 gene family (Chi et al. 2015; Wang et al. 2024b). Based on this updated phylogenetic analysis, we propose several changes that realign nomenclature use to the original proposed clade naming (Synek et al. 2006; Cvrčková et al. 2012). As OsEXO70L members are within the EXO70FX clade, they should retain the original FX designation (Wang et al. 2024b). In addition, as the EXO70J clade is derived from the EXO70B clade, we propose the clade identifier EXO70BX to reflect a similar evolutionary origin and trajectory as observed for the EXO70F/EXO70FX clades.

### EXO70FX is a novel clade that emerged during Poales evolution

The EXO70FX clade is highly expanded and specific to Poales, an extensive order of monocots that includes all agriculturally important cereals. To identify when the EXO70FX clade emerged during Poales evolution, we extracted EXO70 protein sequences from genomes of fourteen representative monocot species, including the recently sequenced genomes of *Ecdeiocolea monostachya* and *Joinvillea ascendens* (‘Ohe) (Takeda-Kimura et al. 2024). Species selected to encapsulate Poales diversity include the basal Poales species *Ananas comosus* (pineapple); members of the Cyperaceae (sedge) and Juncaceae (rush) families; basal graminids; and PACMAD and BOP species within the Poaceae (grass) family (Fig. 2a). *Musa acuminata* (banana) was designated as the outgroup, belonging to the Zingiberales order, which diverged from its sister order Poales 109-123 million years ago (Linder and Rudall 2005).

**Fig. 2.**
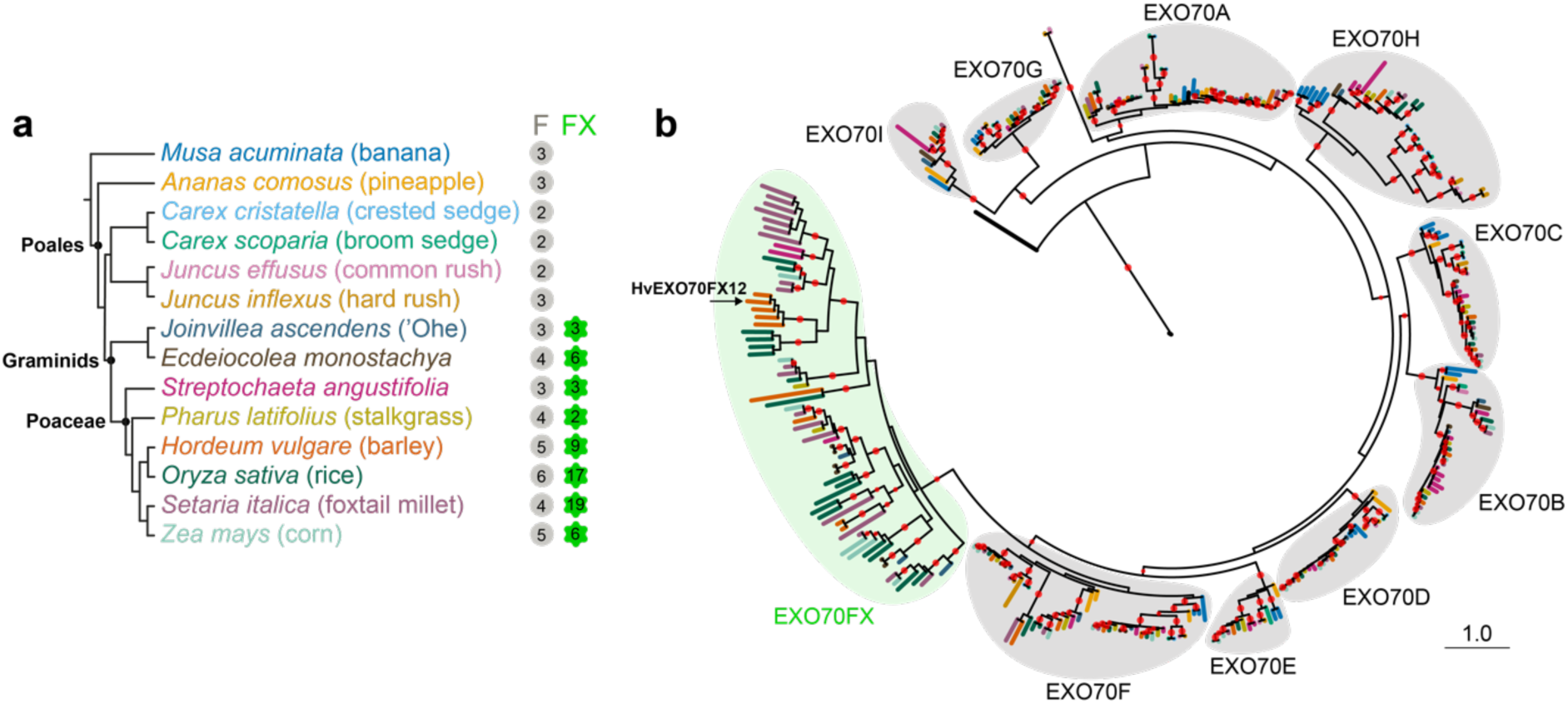
The EXO70FX clade emerged in the Poales after the emergence of sedges and rushes. a) Phylogenetic relationship of 14 monocot species based on previous work, with color-coding corresponding to branches in panel b (Takeda-Kimura et al. 2024). The number of EXO70s belonging to the F and FX clades for each species is noted when present. All evaluated monocot species have members of the F clade, but the FX clade emerges with the emergence of Graminid basal species *Joinvillea ascendens*. b) Structure-based phylogeny of 336 EXO70s from 14 representative monocot species and four EXO70s with solved structures from yeast (PDB 2B1E and 5YFP), mouse (PDB 2PFT), and *A. thaliana* (PDB 4RL5). Red dots indicate bootstrap support greater than or equal to 80%. Scale indicates 1.0 substitution per site.

We performed structure-based alignment and constructed a maximum likelihood phylogenetic tree. Within monocots, EXO70 proteins group into ten clades (A, B, C, D, E, F, FX, G, H, I), all of which are supported by bootstrap support of at least 80%, except for the F clade (Fig. 2b). All clades other than the FX clade are conserved across angiosperms. In contrast, the FX clade appears only with the emergence of *J. ascendens* and is present in all species in the lineage, including *E. monostachya* and all Poaceae species. Within the Poales, the FX clade exhibits the most extreme degree of expansion, the greatest intra-clade divergence, and most variable protein length compared to any other clade (Fig. 2b, Fig. S1). The novelty of the FX clade in graminids, paired with the divergence and expansion of clade members, suggests that the EXO70FX members are fulfilling niche cellular functions either through subfunctionalization or neofunctionalization.

### EXO70FX proteins lack structural requirements for inclusion in the exocyst complex

Several green plant EXO70s have been shown to interact with exocyst subunits and are therefore proposed to function within the exocyst complex. First demonstrated with AtEXO70A1 in plants, exocyst association has been subsequently demonstrated for AtEXO70B1, AtEXO70B2, AtEXO70E2, AtEXO70H1, AtEXO70H4, and OsEXO70H3 (Synek et al. 2006, 2021; Pečenková et al. 2011; Kulich et al. 2013, 2015; Ding et al. 2014; Michalopoulou et al. 2022; Wu et al. 2022). In yeast, initial interactions within the exocyst are mediated by the N-terminal coiled coil CorEx motif of each of the eight subunits (Mei et al. 2018). Within the hierarchal formation of the yeast exocyst, the CorEx domains of EXO70 and EXO84 form an anti-parallel zipper, which then intertwines with the SEC10-SEC15 CorEx zipper (Mei et al. 2018).

To understand the prevalence of the CorEx motif in plant EXO70s, we performed AlphaFold2-based structural predictions of plant EXO70s known to interact with exocyst subunits, yeast and human EXO70s, and HvEXO70FX12 (Fig. S2). Using a structure-based alignment, we predicted the five sub-domains (CorEx, CAT-A, CAT-B, CAT-C, and CAT-D) of each EXO70 based on yeast annotations (Mei et al. 2018; Synek et al. 2021). While the CAT domains were shown to be structurally conserved between all exocyst-interacting EXO70s, the N-terminal region had the greatest divergence (Fig. 3a). The CorEx domains of yeast EXO70, human EXO70, and AtEXO70A1 consist of long rod-like coiled coil CorEx domains. All other plant exocyst-interacting EXO70s contain a predicted coiled coil CorEx domain, but their lengths are highly variable, with members of the EXO70H family having only a short N-terminal coiled coil. In contrast, the predicted structure of HvEXO70FX12 lacks the N-terminal region, and only CAT-B, -C, and -D domains are structurally conserved. Based on the structural divergence of HvEXO70FX12 from exocyst-interacting EXO70s, we predict that HvEXO70FX12 has lost the structural features necessary for its association with the exocyst complex.

**Fig. 3.**
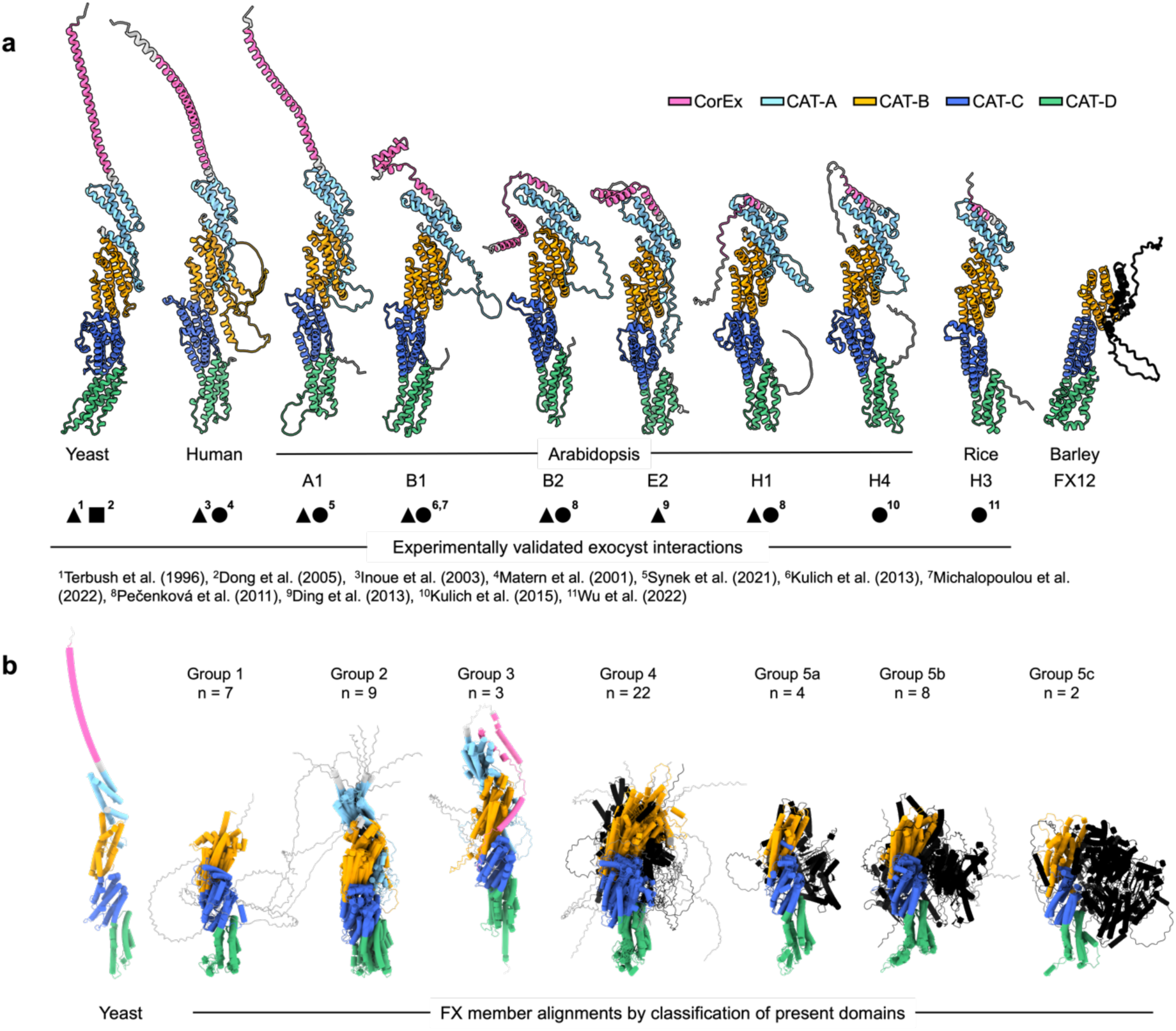
The EXO70FX clade lacks the CorEx domain, which is structurally retained in all exocyst-interacting EXO70s. a) CorEx domains are present in all exocyst-associated plant EXO70s but absent in HvEXO70FX12. The following full-length structures were predicted with AlphaFold2: yeast and human EXO70, plant EXO70s previously shown to interact with exocyst subunits, and HvEXO70FX12. Primary evidence showing exocyst interaction is cited and labelled by shape for the assay performed as follows: pull-downs with purified recombinant proteins, square; any single or combination of co-immunoprecipitation (co-IP), affinity purification followed by mass-spectrometry (AP-MS), bimolecular fluorescence complementation (BiFC), or fluorescence resonance energy transfer (FRET), triangle; and yeast two-hybrid (Y2H), circle. Protein sub-domains from yeast were superimposed on all EXO70s using a structure-based alignment and subsequently colour-coded. b) Predicted structures of EXO70FX proteins from eight monocot species indicate that EXO70FX clade members widely lack the CorEx domain. Structures were predicted with AlphaFold2. Protein sub-domains from yeast were superimposed on all EXO70FX members using a structure-based alignment, curated to ensure structural agreement with yeast, and colour-coded as shown in panel a. EXO70FX members were classified into five groups based on the presence of domains, with the number of proteins identified per group labelled (n). Group members were overlayed and shown with tubular α-helices for clarity.

We next sought to characterize if loss of the CorEx domain was a feature conserved across the EXO70FX clade. We identified 55 EXO70FX members from *J. ascendens*, *E. monostachya*, *H. vulgare*, *O. sativa*, *Z. mays*, *Streptochaeta angustifolia*, *Pharus latifolius*, and *Setaria italica* for structural analysis. Upon defining CorEx and CAT domain boundaries for EXO70FX members based on the structural characterisation of yeast, we observed five main structural groups within the EXO70FX clade based on present domains (Fig. 3b, supplementary table 1). Group 1 contains seven EXO70s that lack both CorEx and CAT-A domains. Group 2 contains nine EXO70s that lack only the CorEx domain, with all other domains conserved. Group 3 contains three EXO70s that have a coiled coil N-terminal of the CAT-A domain, and it is unknown whether these are functional CorEx domains. Group 4, which is the largest group and the one containing HvEXO70FX12, includes 22 EXO70s with a short coiled coil region N-terminal of CAT-B that lacks structural homology with CAT-A. Group 5 contains 14 EXO70s that appear to be fusions with other domains.

Further sub-division of Group 5 found that Group 5a members have a short N-terminal β-sheet domain, Group 5b members have an N-terminal α-β-α fold with homology to the adenine nucleotide alpha hydrolases-like (AANH) superfamily, and Group 5c members are integrated domains within NLRs. Fusion with β-sheet domains was an early acquisition, as Group 5a and Group 5b both emerged with basal graminid species, *E. monostachya* and *J. ascendens*, respectively (Fig. S3). The fusion between an EXO70FX and an AANH domain occurred as a single evolutionary event, which is evidenced with bootstrap support for independent structure-based maximum-likelihood phylogenetic trees for both EXO70FX and AANH domains across the Poales (Fig. S3, Fig. S4). However, integration within NLRs occurred independently in *H. vulgare* and *P. latifolius*, as these EXO70FX domains are more distantly related (Fig. S3, Fig. S5). The divergent N-terminal structures suggest that EXO70FX proteins lost canonical exocyst complex function and have adopted new, unknown function(s).

### HvEXO70FX12 lacks association with exocyst subunits

Previous work in *A. thaliana* has shown interactions between EXO70A1 and exocyst components SEC3A, EXO84B, SEC10, and SEC15B (Synek et al. 2021). To determine if HvEXO70FX12 retains the ability to interact with the exocyst complex, we performed a yeast two-hybrid (Y2H) assay between HvEXO70FX12 and two HvSEC15 paralogs, three HvEXO84 paralogs, and HvSEC3. We found that unlike the positive control AtEXO70A1- AtSEC3A, HvEXO70FX12 did not interact with exocyst subunits, despite all proteins accumulating in yeast (Fig. 4a, Fig. S6).

**Fig. 4.**
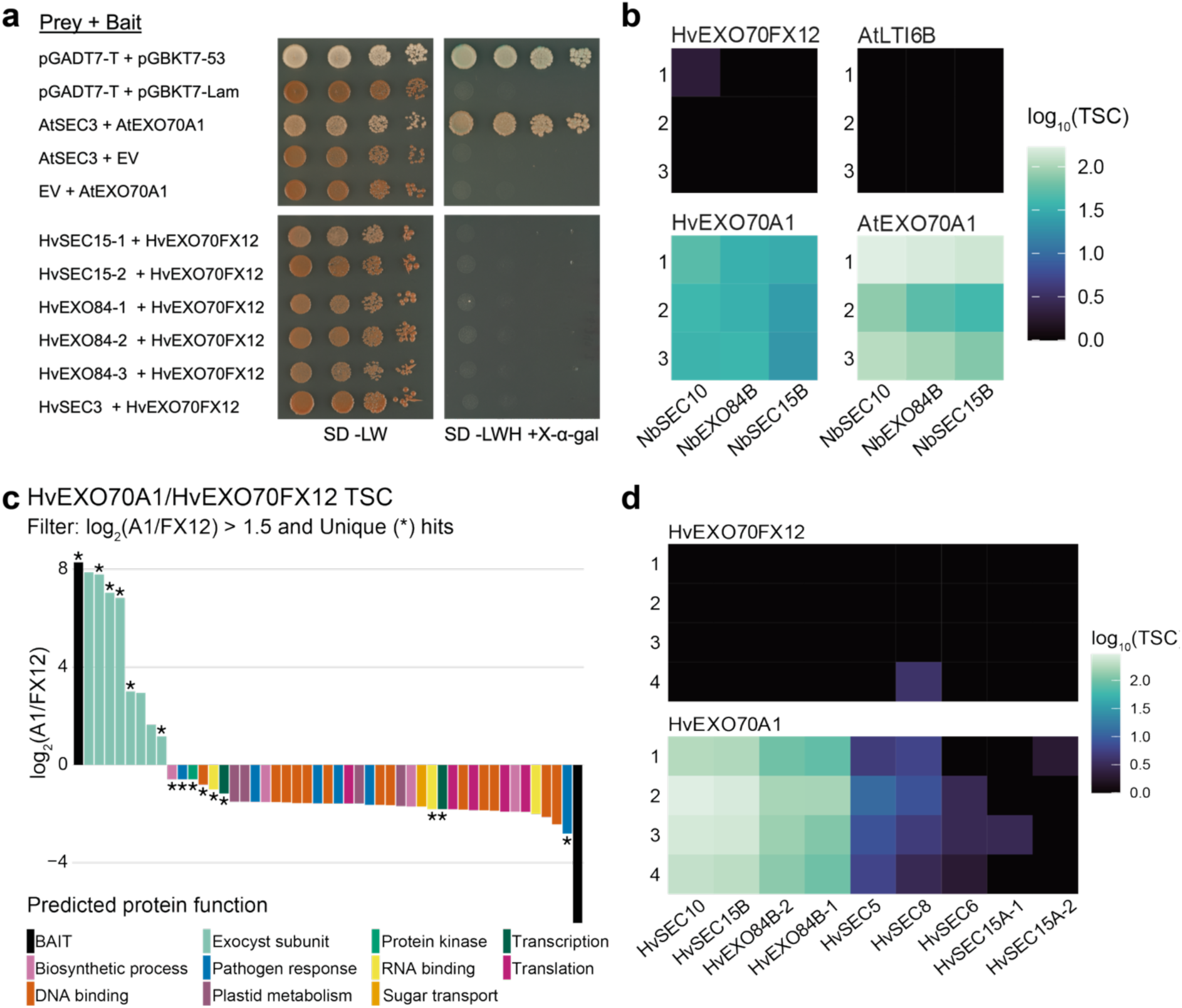
HvEXO70FX12 lacks association with exocyst subunits. a) HvEXO70FX12 does not interact with any exocyst subunit tested in Y2H. Matchmaker® Gold Y2H positive (pGADT7-T + pGBKT7-53) and negative (pGADT7-T + pGBKT7-Lam) controls were used in addition to AtSEC3-AtEXO70A1 as a biologically relevant control (Synek et al. 2021). Growth on synthetic defined (SD) -Leu/-Trp media indicates presence of activation and binding domain plasmids in yeast, while growth on SD/-Leu/-Trp/-His/+X-α-Gal media indicates interaction of bait and prey proteins. Three replicates were performed with similar results. b) HvEXO70FX12 does not associate with any exocyst subunit in *N. benthamiana*. 3xFLAG-HvEXO70FX12, 3xFLAG-HvEXO70A1(positive control), 3xFLAG-AtEXO70A1 (positive control), and AtLTI6B-3xFLAG (negative control) were transiently expressed in *N. benthamiana*. Affinity purification followed by mass spectrometry (AP-MS) was used to identify and quantify associated proteins based on total spectrum count (TSC). HvEXO70A1 and AtEXO70A1 associate with *N. benthamiana* EXO84B, SEC10, and SEC15B proteins. Numbers indicate three independent replicates. c) Unique and enriched barley proteins in HvEXO70A1 pull-down exclusively belong to the exocyst complex. Transgenic barley expressing 3xFLAG-HvEXO70A1 or 3xFLAG-HvEXO70FX12 was used for AP-MS. Associated proteins are shown that are unique (indicated with *) or are at least 2.8X more enriched (log_2_(A1/FX12) > 1.5) based on TSC between samples averaged over four replicates. d) HvEXO70FX12 lacks reproducible association with any barley exocyst subunit in transgenic barley. In contrast, HvEXO70A1 associates with barley SEC10, SEC15B, EXO84B, SEC5B, SEC8, SEC6, and SEC15A proteins. Numbers indicate four independent replicates.

In a complementary approach, we transiently expressed 3×FLAG-HvEXO70FX12, AtLTI6B-3×FLAG, 3×FLAG-HvEXO70A1, and 3×FLAG-AtEXO70A1 in *N. benthamiana*, and performed affinity purification followed by mass spectrometry (AP-MS) to identify associated proteins. As expected, barley and *A. thaliana* EXO70A1 orthologs associated with *N. benthamiana* EXO84, SEC10, and SEC15 paralogs in all three replicates. Strikingly, HvEXO70FX12 and the PM-localised AtLTI6B (negative control) lacked association with all exocyst subunits (Fig. 4b).

Lastly, to exclude the possibility that association with exocyst subunits only occurs in the native context, we created stable barley lines expressing 3×FLAG-HvEXO70A1 or 3×FLAG-HvEXO70FX12 and performed AP-MS. Comparing unique and enriched (≥2.8X) proteins associated with either HvEXO70A1 or HvEXO70FX12 samples indicates that all HvEXO70A1-enriched proteins were exocyst complex members (Fig. 4c). Furthermore, while HvEXO70A1 interacted with SEC10, SEC15B, EXO84B, SEC5B, SEC8, SEC6, and SEC15A, there was a complete lack of reproducible association between HvEXO70FX12 and any exocyst subunit across four replicates (Fig. 4d). Therefore, AP-MS supports the hypothesis that HvEXO70FX12 has lost the ability to function as an exocyst subunit. To preclude the possibility that FLAG tagging impairs the function of HvEXO70FX12, we demonstrated that 3xFLAG-HvEXO70FX12 functionally complements HvEXO70FX12 in *Pst* resistance (Fig. S7).

### HvEXO70FX12 is localised to the PM

To identify HvEXO70FX12 subcellular localisation, we transiently expressed mEGFP- HvEXO70FX12 in *N. benthamiana* via *Agrobacterium tumefaciens*-mediated transformation. HvEXO70FX12 co-localised with the transmembrane chitin receptor AtLYK4 (Fig. 5a). Under conditions of plasmolysis, both mEGFP-HvEXO70FX12 and AtLYK4-Cherry co-migrated away from the cell wall with the PM, suggesting PM rather than apoplastic localisation. To confirm PM localisation within the native context, 35s:mEGFP-HvEXO70FX12 and PM marker 35s:AtLTI6B-mCherry were co-bombarded into barley using biolistic particle bombardment. mEGFP-HvEXO70FX12 localised exclusively to the periphery of the cell and colocalised with AtLTI6B-mCherry, confirming PM localisation (Fig. 5b).

**Fig. 5.**
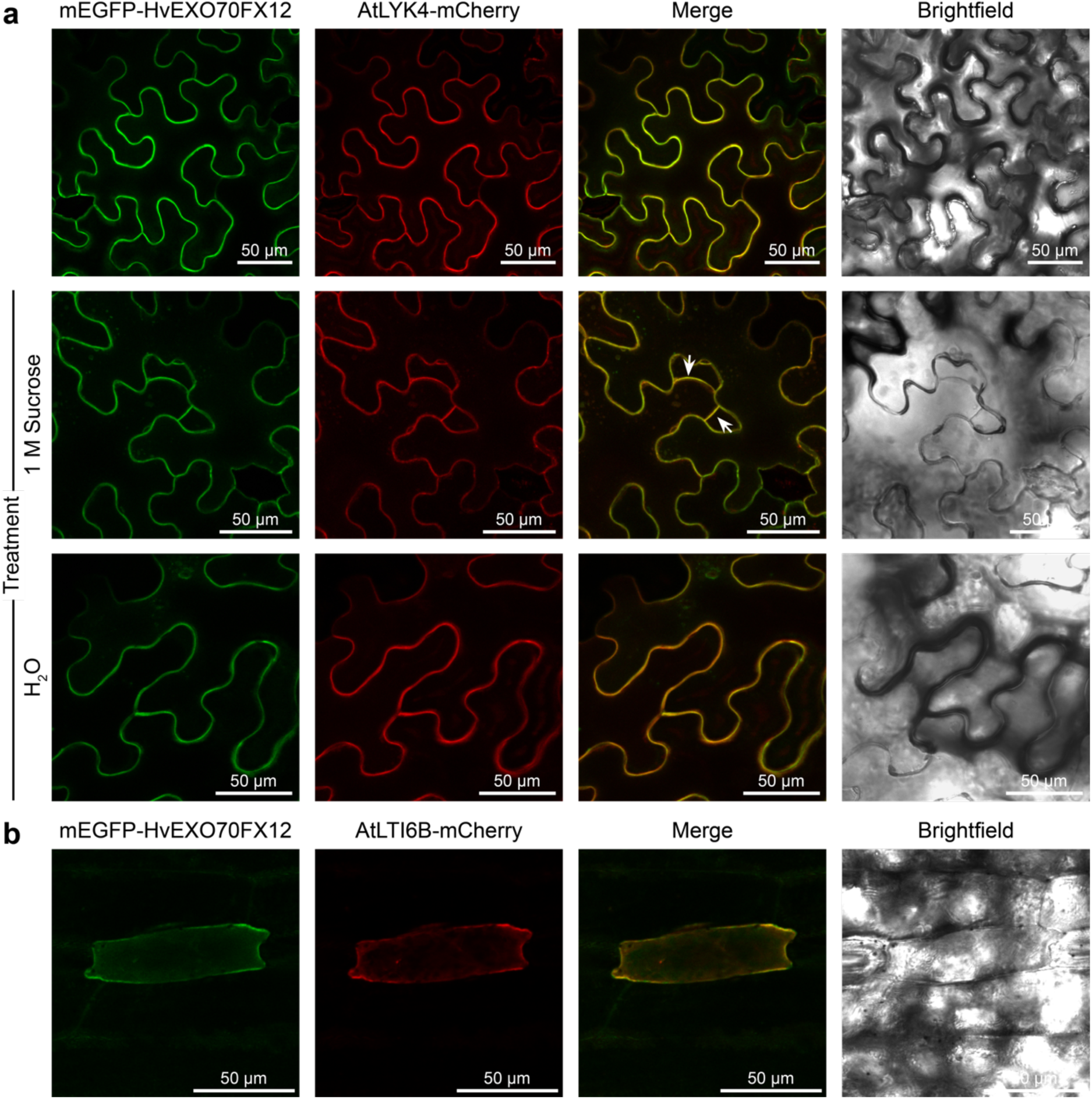
HvEXO70FX12 is plasma membrane (PM)-localised in *N. benthamiana* and barley. a) mEGFP-HvEXO70FX12 co-localises with PM marker AtLYK4-mCherry when transiently co-expressed in *N. benthamiana*. Cells underwent plasmolysis upon treatment with 1.0 M sucrose, and HvEXO70FX12 and AtLYK4 co-localise in the PM during cell shrinkage, as indicated by white arrows. Cells treated with H_2_O as a control did not experience plasmolysis. Two biological replicates were performed with similar results. b) mEGFP-HvEXO70FX12 co-localises with PM marker AtLTI6B-mCherry when transiently co-transformed in barley via biolistic particle bombardment. Two biological replicates were performed with 4-5 bombarded leaves with similar results.

Due to exclusive PM localisation of HvEXO70FX12, we filtered candidate associated proteins from the barley AP-MS dataset based on localisation and included only those predicted to localise to the cytosol or PM (supplementary table 2). Refined candidates include proteins shown to be involved in pathogen responses, namely a remorin, sucrose transporter, Bcl-2- associated athanogene (BAG) domain-containing protein, and ricin B-like lectin, as well as a predicted serine/threonine kinase in the AGC superfamily (Fig. 6).

**Fig. 6.**
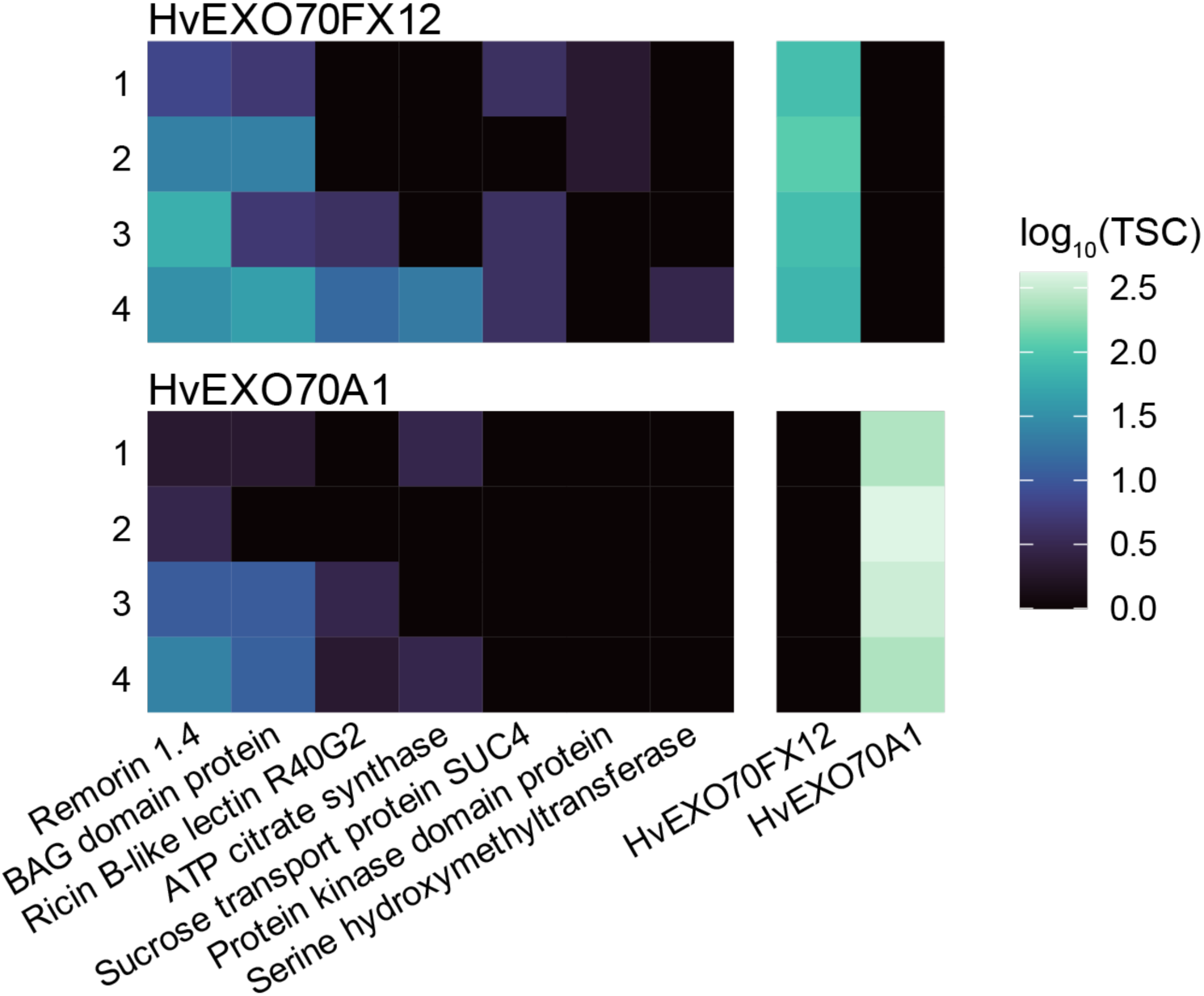
Candidate HvEXO70FX12-associated proteins include defence-related proteins. A heatmap shows differentially enriched proteins between the HvEXO70FX12 and HvEXO70A1 pull-downs from stable barley transgenics based on total spectrum counts (TSC). Candidate genes were first selected based on being unique or ≥2.8X more enriched in the HvEXO70FX12 sample compared to the HvEXO70A1 sample. Candidate proteins were further filtered to only include proteins predicted to localise to the cytosol or PM. Numbers indicate four independent replicates.

## DISCUSSION

EXO70s are uniquely expanded in green plants compared to all other eukaryotic-lineages, and the functional implications of this expansion have remained unresolved. Diverse plant EXO70s have specific functions. While the EXO70A family has been primarily shown to be involved in canonical secretion during plant development, EXO70s from B, E, F, H, and FX families are involved in immunity in various plant-pathogen systems (Pečenková et al. 2011; Stegmann et al. 2012, 2013; Ostertag et al. 2013; Fujisaki et al. 2015; Guo et al. 2018; Wang et al. 2019, 2020; Hou et al. 2020, 202; Holden et al. 2022; Wu et al. 2022; Huebbers et al. 2024). Additionally, EXO70B and EXO70D family proteins have been shown to be involved in autophagy, and the mechanistic connection between autophagy and defence is only beginning to be resolved (Kulich et al. 2013; Acheampong et al. 2020; Brillada et al. 2021). Markovic et al. (2021) demonstrated the specific nature of EXO70 paralogs. Of nine *A. thaliana* EXO70s tested, only EXO70A1 and EXO70A2 could complement *exo70a1* mutant developmental phenotypes, and only EXO70B1, but not EXO70B2, could complement aberrant senescence and anthocyanin accumulation in the *exo70b1* mutant (Marković et al. 2021). HvEXO70FX12 also appears to be highly specific, as related HvEXO70FX11 paralogs in barley cv. Morex are insufficient to confer HvPUR1-mediated resistance to *P. striiformis* f. sp. *tritici* (Holden et al. 2022). HvEXO70FX11b has been shown to positively regulate resistance in barley to another fungal pathogen: barley powdery mildew (*Blumeria graminis* f. sp. *hordei*) (Ostertag et al. 2013).

Multiple hypotheses have been proposed to describe how EXO70 diversification applies to exocyst function. First, EXO70s may dictate specific forms of the exocyst with distinct functions. For example, subfunctionalization can occur in which EXO70 paralogs have specific functions as exocyst subunits due to distinct tissue-specific expression or subcellular localisations. When expressed under the *AtEXO70A1* promoter in an *exo70a1* mutant background, all paralogs tested showed punctate localisation in *A. thaliana* roots except for the PM-localising AtEXO70A1 and AtEXO70A2 (Marković et al. 2021). Similarly, under the *AtEXO70H4* promoter in a *exo70h4* mutant background, AtEXO70A1 and AtEXO70B1 had predominantly cytosolic accumulation in trichomes rather than mimicking the trichome PM localisation of AtEXO70H4, suggesting cell-type specific localisations (Kulich et al. 2018). As AtEXO70A1 and AtEXO70H4 secrete specific cargo to the PM, such as PIN auxin efflux carriers and callose synthases, respectively (Drdová et al. 2013; Kulich et al. 2018), paralog-dependent localisation enables tightly regulated exocytosis to cells that require a specific and dynamic response. Alternatively, EXO70 paralog specificity may dictate the inclusion of specific exocyst subunits, leading to distinct exocyst forms. For example, the exocyst-positive organelle (EXPO) is a proposed double-membrane exocytotic structure in *A. thaliana* that is mediated by EXO70E2 and contains SEC5A, SEC6, SEC8, and SEC10, but not EXO84B (Ding et al. 2014). EXPOs, which localise in cytosolic puncta, are thought to be involved in unconventional protein secretion (Ding et al. 2014).

It has long been suggested that EXO70 expansion has enabled EXO70 paralogs to act independently of the exocyst complex. Under the balance hypothesis, an imbalance of one subunit within a complex could have deleterious effects on the complex and thus is under negative selection (Papp et al. 2003; Synek et al. 2006). Within green plants, the expansion of EXO70s is unique, with SEC3, SEC5 SEC6, SEC8, and SEC10 generally having only one copy and SEC15 and EXO84 generally having two to four copies (Cvrčková et al. 2012). While it has not been shown how copy number of EXO70s affects accumulation of EXO70s compared to other exocyst subunits in cell-type specific contexts, the extreme expansion of EXO70s is at odds with the equal stoichiometries of other exocyst subunits (Cvrčková et al. 2012). Recently, it has been shown that *Marchantia polymorpha* EXO70II has reduced capacity to associate with the exocyst complex due to a negatively charged and structurally divergent N-terminal region (De La Concepcion et al. 2024). As *M. polymorpha* EXO70II is the first basal member to be functionally characterized in the EXO70.2 clade, encompassing angiosperm EXO70B, EXO70C, EXO70D, EXO70E, EXO70F, EXO70FX, and EXO70H clades, an initial hypothesis is that these families either retain a full or subset of functions from the common ancestor of the EXO70.2 clade. Interestingly, charge status of the N-terminal is not preserved in the EXO70.2 clade, with the EXO70H family retaining a positive charge (De La Concepcion et al. 2024). In addition, several molecular and biochemical studies have found exocyst complex association for EXO70 family members in the EXO70B and EXO70H clades (Pečenková et al. 2011; Kulich et al. 2013, 2015; Michalopoulou et al. 2022). Therefore, it remains unclear if *M. polymorpha* EXO70II represents ancestral function for the EXO70.2 clade or has experienced a unique evolutionary trajectory in an early branching lineage. We similarly observe that HvEXO70FX12 functions independently from the exocyst complex, although this appears to be a recent innovation through the rapid loss of CorEx and CAT-A domain loss. Based on the CorEx domain loss in 52 of 55 EXO70FX proteins, we predict that this Poales-specific clade has widely undergone neofunctionalization.

In yeast and mammals, EXO70 has been suggested to have exocyst-independent functions not demonstrated in plants. Yeast and rat EXO70 interact with ARPC1, a subunit of the Arp2/3 complex that regulates actin reorganization for cell motility (Zuo et al. 2006). EXO70 promotes the interaction between Arp2/3 and the nucleation promoting factor (NPF) WAVE2, which leads to enhanced actin filament nucleation and branching (Liu et al. 2012). While the exocyst complex may interact with Arp2/3, EXO70 was sufficient in isolation to stimulate actin polymerization *in vitro* (Liu et al. 2012; Zhu et al. 2019). Additionally, rat EXO70 impacts cell shape and migration by forming oligomers and creating negative membrane curvature (Zhao et al. 2013).

Determining the exocyst-independent nature of HvEXO70FX12 enables further unbiased exploration of the functional role of HvEXO70FX12 in wheat stripe rust resistance and more broadly, the role of the EXO70FX family. Due to the reciprocal dependency of HvEXO70FX12 and HvPUR1 in resistance, we postulate that HvEXO70FX12 functions within this LRR-RK-triggered pathway, with LRR-RKs closely related to HvPUR1, including AtFLS2, AtEFR, and OsXA21, having well-established roles as PRRs in eliciting PTI upon extracellular perception of microbial patterns (Song et al. 1995; Gómez-Gómez and Boller 2000; Zipfel et al. 2006). Surprisingly, no interaction between HvEXO70FX12 and HvPUR1 has been detected from Y2H and proteomics approaches (Fig. S8), so the mechanistic relationship of HvEXO70FX12 in HvPUR1-mediated signalling remains unclear.

We hypothesize that HvEXO70FX12 is involved in PTI through candidate associated proteins. Several defence-related proteins enriched in the HvEXO70FX12 pull-down in transgenic barley offer initial insight into candidate immune responses that involve HvEXO70FX12 in the native context. Preliminary evidence suggests that HvEXO70FX12 associates with a remorin, which belongs to a plant-specific family of PM-anchored proteins (Raffaele et al. 2007). Remorins have been implicated in diverse immune responses, including stabilizing membranes (Legrand et al. 2023), engineering PM nanodomains enriched in RKs and PRR signalling complexes (Bücherl et al. 2017; Liang et al. 2018; Traeger et al. 2023; Wang et al. 2024a), regulating cell-to-cell movement through plasmodesmata conductance (Perraki et al. 2018; Rocher et al. 2022), and enhancing cell death (Bozkurt et al. 2014; Cai et al. 2020). Additional putative HvEXO70FX12-associated proteins include a sucrose transporter, belonging to a protein family implicated in sugar compartmentalization between plants and biotrophic pathogens (Sun et al. 2010; Moore et al. 2015; Liu et al. 2022) and a BAG domain-containing protein, belonging to a protein family that has been implicated in regulating diverse stress and developmental responses, including fungal resistance in *A. thaliana* and rice (Kabbage and Dickman 2008; Li et al. 2016; You et al. 2016).

The pattern of evolutionary diversification and expansion observed in the EXO70FX clade aligns with that of NLRs and PRRs in plants and supports an alternative hypothesis that members of the EXO70FX may be involved in sensing pathogens (Lehti-Shiu et al. 2012; Ngou et al. 2022). The possibility cannot be excluded that some EXO70FX members act as decoys by interacting with effectors, only to have these interactions monitored by guard proteins that execute immune signalling, or some EXO70FX members may fulfil helper roles by interacting with and activating effectors for host recognition (Dangl and Jones 2001; Van Der Hoorn and Kamoun 2008; Win et al. 2012). In this study, we identified two EXO70FX members that are integrated domains in NLRs from barley and stalkgrass (*P. latifolius*). EXO70F1 has previously been shown to be integrated into the NLR HvRGH2 prior to the Poeae-Triticeae radiation in grass evolution, indicating that NLR integration is not specific to the FX clade (Brabham et al. 2018). While not an integrated domain, OsEXO70F3 interacts with AVR-Pii and is required for rice blast resistance conferred by the NLR pair OsPII-1 and OsPII-2, likely through a helper or guard model (Fujisaki et al. 2015). As AtEXO70B1 is the target of multiple bacterial effectors that promote susceptibility, the diversity of EXO70s in plants likely led to the evolution of decoys, whether they are integrated in NLRs or interacting with NLRs to perceive and respond to effectors. Interestingly, eight EXO70FX members are fused to AANH- like domains, and each is derived from a single fusion event that occurred during Poales evolution. The AANH superfamily is a family of proteins with an ATP-binding α-β-α fold that includes diverse enzymes involved in primary metabolism and sulphur transferases (Litomska et al. 2018). The EXO70FX-fused AANH domains are derived from class VI plant U-box proteins (PUBs), which are a family of U-box-protein kinase fusion proteins (Trenner et al. 2022). It is unclear why some EXO70FX members are fused to this domain and whether these EXO70 domains may be serving as decoys, enzymatic, or signalling roles.

In conclusion, we show that a barley EXO70 has lost association with the exocyst complex. The EXO70FX clade is a novel acquisition in grasses and grass-like species that has undergone diversification, expansion, and likely neofunctionalization. We predict that EXO70FX members have lost the ability to interact with the exocyst and may have diverse mechanisms in immune pathways. Determining the role of HvEXO70FX12 within the HvPUR1-mediated defence pathway and the greater role EXO70FX members play in immunity will shed light on the evolution of a lineage-specific feature of plant immunity.

## METHODS

### Phylogenetic Analysis

Publicly available genomes were retrieved for 23 green plant species used in the green plant analysis, 14 monocot species including for EXO70FX clade emergence analysis, and 14 overlapping monocot species for AANH analysis (supplementary dataset 1). We identified PFAM protein domains for all proteins with InterProScan and selected proteins annotated with the EXO70 domain (PF03081) or the AANH domain (SUPERFAMILY identifier SSF52402). Structure-based alignments were performed with MAFFT-DASH (v7.520) based on protein structures deposited in the Protein Data Bank (PDB). Alignments were filtered to remove identical proteins, filtered to only include proteins with 40% sequence coverage and residues with 20-40% coverage, and curated manually. Trees were constructed using RAxML (v8.2.12) with GAMMA model of heterogeneity, an automatically optimised substitution matrix, and 1,000 bootstraps. Full-length proteins were used in the green plant analysis, while the EXO70FX clade emergence analysis and AANH domain analysis were performed with extracted domains only.

### Structural Predictions

To compare EXO70s in plants, yeast, and human for subdomain analysis, structures were predicted using ColabFold v1.5.2-v1.5.5 AlphaFold2 with default settings. The following sequences were used for structure prediction: AtEXO70A1 (AT5G03540.1), AtEXO70B1 (AT5G58430.1), AtEXO70B2 (AT1G07000.1), AtEXO70E2 (AT5G61010.1), AtEXO70H1 (AT3G55150.1), AtEXO70H4 (AT3G09520.1), OsEXO70H3 (LOC_Os12g01040), HvEXO70FX12 (HORVU.MOREX.r3.4HG0407730.1), *S. cerevisiae* EXO70 (UniProt P19658) and *Homo sapiens* EXO70 (UniProt Q9UPT5). The AlphaFold2-predicted structures of ScEXO70 and AtEXO70A1 were used for visualisation of domains rather than the available crystal structures due to truncation of the N-terminal rod-like structures in crystal structures. To ensure highly accurate predictions, the predicted structures of ScEXO70 and AtEXO70A1 were overlayed on crystal structures from the PDB (ScEXO70: 2B1E and 2B7M; AtEXO70A1: 4RL5) in ChimeraX1.8 with Matchmaker default settings, and excellent chain pairing was confirmed.

For the EXO70FX dataset, structures were predicted with AlphaFold2 (v2.3.1) with monomer models and PDB selection date threshold of 2023-07-26. A list of EXO70FX members used in subdomain analysis is provided (supplementary table 1).

To superimpose yeast domains, a MAFFT alignment was created using plant EXO70s of interest in addition to EXO70s from diverse eukaryotic lineages used in domain analysis by Synek et al., 2021: *Coccomyxa subellipsoidea* (UniProt I0YQJ0), *Mus musculus* (UniProt O35250), *Physcomitrium patens* (UniProt A0A7I4A6I0), *Drosophila melanogaster* (UniProt Q9VSJ8), *Caenorhabditis elegans* (UniProt P91149), *Dictyostelium discoideum* (UniProt Q558Z9), *Trypanosoma brucei* (UniProt A0A3L6L1H3), and *Schizosaccharomyces pombe* (UniProt Q10339). Yeast domain boundaries were superimposed onto EXO70s based on the MAFFT structure-based alignment, and boundaries were manually curated by aligning EXO70s of interest to ScEXO70 in ChimeraX1.8 with the Matchmaker default settings.

### Plasmid Construction

For Y2H assays, coding sequences of the following genes were digested and ligated into pGADT7 and pGBKT7 vectors provided by TSL SynBio via Golden Gate Assembly: *AtSec3* (AT1G47550.2, TSL SynBio pICSL80078), *AtExo70A1*, *HvSec15-1* (HORVU.MOREX.r3.3HG0259220.1), *HvSec15-2* (HORVU.MOREX.r3.2HG0197180.2), *HvExo84-1* (HORVU.MOREX.r3.5HG0522090.1), *HvExo84-2* (HORVU.MOREX.r3.2HG0133880.1), *HvExo84-3* (HORVU.MOREX.r3.2HG0145850.2), *HvSec3* (HORVU.MOREX.r3.4HG0341340.1), and *HvExo70FX12*.

For proteomics analysis of transiently transformed *N. benthamiana*, *AtLTI6B* (AT3G05890.1^C26F^, TSL SynBio pICSL80100), *HvExo70A1* (HORVU.MOREX.r3.2HG0213760.1), *AtExo70A1*, and *HvExo70FX12* were each fused to a *3×Flag* tag (TSL SynBio pICSL30005/pICSL50007) and assembled into the following level 1 expression cassettes via Golden Gate Assembly: *Act2:LTI6B-3xFlag:Ocs*, *Ubi10:3×Flag-HvExo70A1:Ocs*, *Ubi10:3×Flag-AtExo70A1:Ocs*, and *Ubi10:3×Flag-HvExo70FX12:Ocs*. The ubiquitin and actin promoters were derived from *A. thaliana* (TSL SynBio pICSL12015/pICSL13005; pICH87644), and the octopine synthase terminator was derived from *A. tumefaciens* (TSL SynBio pICH41432).

Level 2 three-gene expression cassettes were created for stable transformation in the barley accession SxGP DH-47, which lacks *HvExo70FX12* and *HvPur1* (Fig. S9). Plasmids were created via Golden Gate Assembly in the acceptor plasmid pICSL4723 (TSL SynBio), which expresses kanamycin resistance conferred by *NptII* in the backbone. All level 2 transgenics included a hygromycin selection marker conferred by *HptII* (TSL SynBio pICSL80036) in reverse orientation under control of the CaMV+TMV *35s* promoter (TSL SynBio pICH51277) and *AtHSP18* terminator (TSL SynBio pICSL60008). Transgenic lines additionally overexpressed *HvPur1* (HORVU.MOREX.r3.4HG0407750.1) under the control of the *ZmUbi* promoter (TSL SynBio pICSL12009) and the CaMV *35s* terminator (TSL SynBio pICH41414). For lines used in barley proteomics experiments, *HvPur1* was tagged with *4×Myc* (TSL SynBio pICSL30009), which was embedded in the N-terminal region immediately preceding the first LRR motif between HvPUR1^73Q^ and HvPUR1^74V^. For the transgenic line created to confirm *3×Flag-HvExo70FX12* function, *HvPur1* was untagged (Fig. S9). The third gene in the transgenic cassettes consisted of *HvExo70A1* or *HvExo70FX12*. Both were tagged N-terminally with *3×Flag* (TSL SynBio pICSL30005) and under control of the *OsAct1* promoter (TSL SynBio pICSL13017) and nopaline synthase (*Nos*) terminator (TSL SynBio pICH41421) derived from *A. tumefaciens*.

For subcellular localisation in *N. benthamiana, Ubi10:mEGFP-HvExo70FX12:Ocs* was generated in the expression vector pICH47732 (TSL Synbio) with an mEGFP N-terminal tag (TSL SynBio pICSL30032). The *35s:3×HA-AtLyk4-mCherry:HSP18* construct was generated in the expression vector pICSL86900_OD (TSL SynBio), using golden gate cloning by combining the level 0 parts of the *35s* promoter (TSL Synbio pICH41388), the first 21 AA of the AtLYK4 coding sequence as the *AtLyk4* signal peptide in pICH41246 (TSL Synbio), an N-terminal *3×HA* tag (ENSA EC15191), the coding sequence of *AtLyk4* without the signal peptide sequence in pAGM1299 (TSL Synbio), a C-terminal mCherry tag (TSL Synbio pICSL50004), and the terminator of the *AtHSP18* gene (ENSA EC15320). For subcellular localisation in barley, *35s:mEGFP-HvExo70FX12:35s* and *35s:LTI6B-mCherry:HSP18* were generated in the expression vector pICH47732 (TSL Synbio *p35s*: pICSL13002/pICH51277; *mEGFP*: pICSL30032; mCherry: pICSL50004; *t35s*: pICH41414; *tHSP18*: pICSL60008).

### Transient Expression in *N. benthamiana*

Plasmids were transformed into *A. tumefaciens* strain GV3101 with electroporation. Liquid cultures of *A. tumefaciens* carrying the desired plasmids were incubated overnight in LB medium at 28°C with constant shaking. Bacteria was pelleted with centrifugation (1290 rcf) for 10 min and resuspended in infiltration buffer (10 mM MES pH 5.6, 10 mM MgCl_2_, and 150 μM acetosyringone). Bacterial suspensions were diluted to an OD_600_ of 0.2 for AP-MS experiments and OD_600_ of 0.4 for microscopy experiments prior to infiltration into expanded leaves of four-week-old *N. benthamiana* plants. Samples were collected 2-3 days after infiltration.

### Y2H Assays

Gold Yeast cells were made competent chemically and transformed with bait and prey plasmids according to the Frozen-EZ Yeast Transformation II™ protocol. Transformed cells were grown in liquid synthetic defined (SD) medium that lacked leucine and tryptophan for approximately 1 hr, plated on SD/-Leu/-Trp medium, and incubated at 27°C for approximately four days. One colony was selected for each prey/bait pair and inoculated in SD/-Leu/-Trp overnight. Yeast cultures were plated on SD/-Leu/-Trp and SD/-Leu/-Trp/-His/+X-α-Gal in four serial dilutions starting at OD of 1.0 with each sequential dilution being 1:100 of the former. Growth on SD/-Leu/-Trp confirmed yeast transformation with both the pGBKT7 plasmid encoding the bait and a tryptophan biosynthesis gene and the pGADT7 plasmid encoding the prey and a leucine biosynthesis gene. Growth on SD/-Leu/-Trp/-His/+X-α-Gal confirmed interaction of transformed proteins, as physical proximity of the GAL4 binding domain and activation domain activates transcription of reporter genes including a histidine biosynthesis gene and an a-galactosidase enzyme that turns yeast colonies blue in the presence of X-α-Gal. A Matchmaker® Gold Y2H positive control of pGADT7-T + pGBKT7-53 (T-antigen and murine p53) and negative control of pGADT7-T + pGBKT7-Lam (T-antigen and lamin) controls were used in addition to biological controls. Colonies were incubated at 27°C for approximately four to six days and then photographed.

We followed a protocol adapted from De la Concepcion et al. to detect protein accumulation in yeast via immunoblotting (De La Concepcion et al. 2021). Specifically, 2 mL of yeast cells resuspended in ddH_2_O were pelleted and resuspended in 100-200 µL 4× Laemmli Buffer. The solution was boiled at 95°C for 15 min and centrifuged for 2 min at 800 rpm prior to electrophoresis and transfer to a membrane. Membranes were incubated in blocking solution of 5% milk in TBST (TBS, 0.1% Tween). Proteins were detected with the anti-GAL4 BD (Sigma G3042) or anti-GAL4 AD (Sigma G9293) primary antibodies at 1:5,000 concentration followed by an anti-rabbit secondary antibody (Sigma A0545) at 1:30,000. Each blocking or antibody incubation step occurred for 1 hr at approximately 22°C or overnight at 4°C with constant shaking. Four membrane washes were performed before and after secondary antibody incubation with TBST, and the final wash prior to detection was performed with TBS. Membranes were treated with ThermoScientific™ Femto developing solution prior to chemiluminescence detection in an Amersham ImageQuant 800 (Cytiva) imaging system. Membranes were stained with Ponceau Red to verify equal protein loading across samples.

### Transgenic barley materials

*A. tumefaciens* strain AGL1 was transformed with plasmid DNA via electroporation and recovered in 700 µL L medium at 28°C in constant movement for 1 hr. Transgenic cassettes were then transformed into *H. vulgare* cv. SxGP DH-47 via *Agrobacterium*-mediated transformation according to the protocol described by Hensel et al. (Hensel et al. 2009). Single copy transgenic insert T_1_ lines were selected based on copy number analysis performed by AttoDNA (Norwich, UK). Three independent segregating T_1_ families with high accumulation of 3×FLAG-HvEXO70FX12 or 3×FLAG-HvEXO70A1 were selected for transgenic barley AP-MS replicates. Standard growth conditions of transgenic barley consisted of 18°C 16-hr day/12°C 8-hr night.

### AP-MS

For transient expression in *N. benthamiana*, 4-6 leaves were collected 2-3 days after infiltration. For stable expression in barley, first leaves from approximately 16 plants per T_1_ family were collected 12-15 days after sowing. Plant tissue was snap frozen in liquid nitrogen and ground by mortar and pestle. Double the volume of extraction buffer as plant tissue was added to the tissue (50 mM Tris pH 7.5, 150 mM NaCl, 2.5 mM EDTA, 10% glycerol, 1% IGEPAL CA-630, 5 mM DTT, and 1% plant protease inhibitor (Sigma 9599)). Protein was solubilized by rotating at 4°C for 30 min. Protein was filtered through miracloth and centrifuged (4°C, 30000×g). Input samples were collected before incubating the remaining sample with Anti-FLAG M2 Affinity Gel (Sigma-Aldrich) beads for 2 hr at 4°C. Samples were centrifuged and washed three times with wash buffer (50 mM Tris pH 7.5, 150 mM NaCl, 2.5 mM EDTA, 10% glycerol, 0.5% IGEPAL CA-630). After the final washing step, samples were boiled in 4× NuPage Buffer (70°C, 10 min), centrifuged, and loaded to a pre-cast NuPage gel. Samples were run on the gel at 120 V for 15-30 min. Samples were stained with the SimplyBlue™ Safe Stain (ThermoFisher Scientific) and washed according to the manufacturer directions.

In-gel affinity-enriched proteins were digested by trypsin after reduction by DTT and carbamidomethylation by chloroacetamide. Extracted peptides were measured by liquid chromatography coupled to mass spectrometer Orbitrap Fusion (Thermo), a tandem mass spectrometer operated in data-dependent acquisition mode, or to mass spectrometer timsTOF Pro (Bruker), operated in positive PASEF mode. Raw files were peak-picked by MSConvert (Proteowizard), searched by Mascot (Matrix Science Ltd) against peptide sequences defined by the *H. vulgare* cv. Morex v3 genome (Phytozome ID: 702; Mascher et al. 2021) or *N. benthamiana* transcriptome assembly v.6.1 (https://www.nbenth.com, Queensland University of Technology), and loaded to Scaffold v.5.3.3. (Proteome Software Inc). Based on the evaluation of assigned decoys in a Percolator probability distribution for each dataset, the *N. benthamiana* total spectrum count (TSC) dataset was filtered and exported with a 99.9% protein threshold and 80% peptide threshold, and the barley TSC dataset was filtered and exported with 1% false discovery rate (FDR). Missing values were imputed as 1 to enable log fold-change calculations. For a pairwise comparison between HvEXO70FX12 and HvEXO70A1 enriched proteomes, hits were filtered to include only those found in at least two reps that were unique to A1 or FX samples or enriched with a log_2_(A1/FX12) fold change greater than 1.5. Heatmaps show exocyst subunit hits from *N. benthamiana* and barley datasets: NbSEC10 (Nbv6.1trP38996), NbSEC15B (Nbv6.1trP32490), NbEXO84B (Nbv6.1trP33967), HvSEC5B (HORVU.MOREX.r3.2HG0167650.1), HvSEC6 (HORVU.MOREX.r3.6HG0614830.1), HvSEC8 (HORVU.MOREX.r3.7HG0691530.1), HvSEC10 (HORVU.MOREX.r3.5HG0453940.1), HvSEC15A-1 (HORVU.MOREX.r3.4HG0402500.1), HvSEC15A-2 (HORVU.MOREX.r3.2HG0197180.2), HvSEC15B (HORVU.MOREX.r3.3HG0259220.1), HvEXO84B-1 (HORVU.MOREX.r3.5HG0522090.1), and HvEXO84B-2 (HORVU.MOREX.r3.2HG0145850.2). To assess HvEXO70FX12-associated candidate proteins, proteins were filtered by localisation based on subcellular compartment predictions with Panther19.0 (https://pantherdb.org/) and DeepTMHMM (Hallgren et al. 2022).

### Particle bombardment

Barley was grown at a 18°C 16-hr day/16°C 8-hr night regime for seven days. Samples of approximately 1 cm x 1.5 cm were cut from the tips of first leaves and suspended on 0.8% water agar. High-concentration plasmid DNA was prepared with the QIAGEN® Plasmid *Plus* Midi Kit according to manufacturer instructions. Gold microcarriers were prepared and loaded with 4 µg of each plasmid DNA as described by Tee et al. (Tee et al. 2022). DNA-coated gold particles were bombarded into the abaxial side of leaf samples under 1100 PSI with a PDS- 1000/He Particle Delivery System (Bio-Rad). Transformed leaves were incubated abaxial-side down on moistened 0.8% w/v water agar plates in the dark at 4°C for 2-3 days prior to imaging.

### Confocal microscopy

*N. benthamiana* and barley leaf samples were imaged with a Leica SP8X confocal microscope with line sequential scanning. The fluorophore tags mEGFP and mCherry were excited using a white light laser (WLL) with wavelengths at 488 nm and 587 nm and detected at 498-550 nm and 592-640 nm, respectively. For barley specimens, minor gating (beginning after 0.1-0.2 ns) was performed to reduce autofluorescence. For plasmolysis, *N. benthamiana* leaf disks were floated in 1.0 M sucrose or water for 30 min prior to imaging.

### Barley-wheat stripe rust infection

Pathogen assays were conducted as previously described in detail by Holden et al. (Holden et al. 2022). Briefly, seedlings were grown in John Innes peat-based compost at 18°C 16-hr day/12°C 8-hr night, inoculated with *Puccinia striiforms* f. sp. *tritici* isolate 20/092 at 10 days after sowing via pressure spraying in a talcum powder suspension, incubated within a moist, dark sealed box at 4°C for 48 hr, and subsequently grown at 18°C 16-hr day/12°C 8-hr night. First leaves were scored 14 days after infection for pustule development as described by Dawson et al. (Dawson et al. 2015). The scale represents the percentage of the leaf surface that is covered with pustules from none (0) to 100% (4) with intervals of 0.5 based on what is visibly discernible. To compare infection scores between genotypes in pathogen assays, we performed a Wilcoxon-Mann-Whitney (WMW) test (α ≤ 0.05) because datasets did not have normal distributions or homogenous variance between groups.

## Acknowledgements

We are very thankful to Samuel Holden, who laid the groundwork for this project. We thank Brian Steffenson, Dan Voytas, and Colby Starker for their insight and support. We also thank TSL SynBio, TSL Media Kitchen, Kostya Kanyuka, Adam Donaldson, Phil Robinson, and Janina Tamborski for their contributions. The authors acknowledge the Minnesota Supercomputing Institute (MSI) at the University of Minnesota for providing resources that contributed to the research results reported within this paper (http://www.msi.umn.edu). This research used resources provided by the SCINet project and/or the AI Center of Excellence of the USDA Agricultural Research Service, ARS project numbers 0201-88888-003-000D and 0201-88888-002-000D.

## Grants

Fulbright US Student Award (2020-2021) supporting MB

The Gatsby Charitable Foundation, supporting MJM and CZ

UKRI-BBSRC grant APH-ISP BB/X010996/1, supporting CF, MJM

United States Department of Agriculture award number 58-5062-1-006, supporting HM

United States Department of Agriculture-Agricultural Research Service CRIS #5062-21220-025-000D, supporting MJM

## Data availability

Sequencing data used in this study are found in the NCBI database under BioProject codes PRJNA761551, PRJNA378723, and PRJNA1138684. The mass spectrometry proteomics data have been deposited to the ProteomeXchange Consortium via the PRIDE partner repository with the dataset identifier PXD057065 and PXD057109 for *N. benthamiana* and barley datasets, respectively. All other data are available in the main text or the supplementary materials. Raw data, uncropped images, FASTA files, multiple sequence alignments, phylogenetic trees, and scripts used for data analysis and figure preparation are available on Ag Data Commons (https://doi.org/10.15482/USDA.ADC/27997025). A material transfer agreement with The Sainsbury Laboratory is required to receive plasmids. The use of the materials will be limited to non-commercial research uses only. Please contact M.J.M. regarding biological materials, and requests will be responded to within 60 days.

## Author contributions

Conceptualization: M.B., J.R., C.F., M.J.M.

Methodology: M.B., J.S., M.J.M.

Validation: M.B., J.S., M.J.M.

Formal analysis: M.B., J.S., M.J.M.

Investigation: M.B., J.S., I.H-P., P.G, J.T., M.S., S.S., M.A., A.T., H.M.

Resources: C.Z, C.F., M.J.M

Data Curation: M.B., J.S., M.J.M.

Writing - Original Draft: M.B.

Writing - Review & Editing: M.B., S.S., J.R., H.M., J.S., F.L.H.M., C.F., C.Z., M.J.M.

Visualization: M.B. and M.J.M.

Supervision: C.Z., J.R., C.F., and M.J.M.

Project administration: M.J.M.

Funding acquisition: M.B., C.Z., and M.J.M.

## SUPPLEMENTARY MATERIAL

**Fig. S1.**
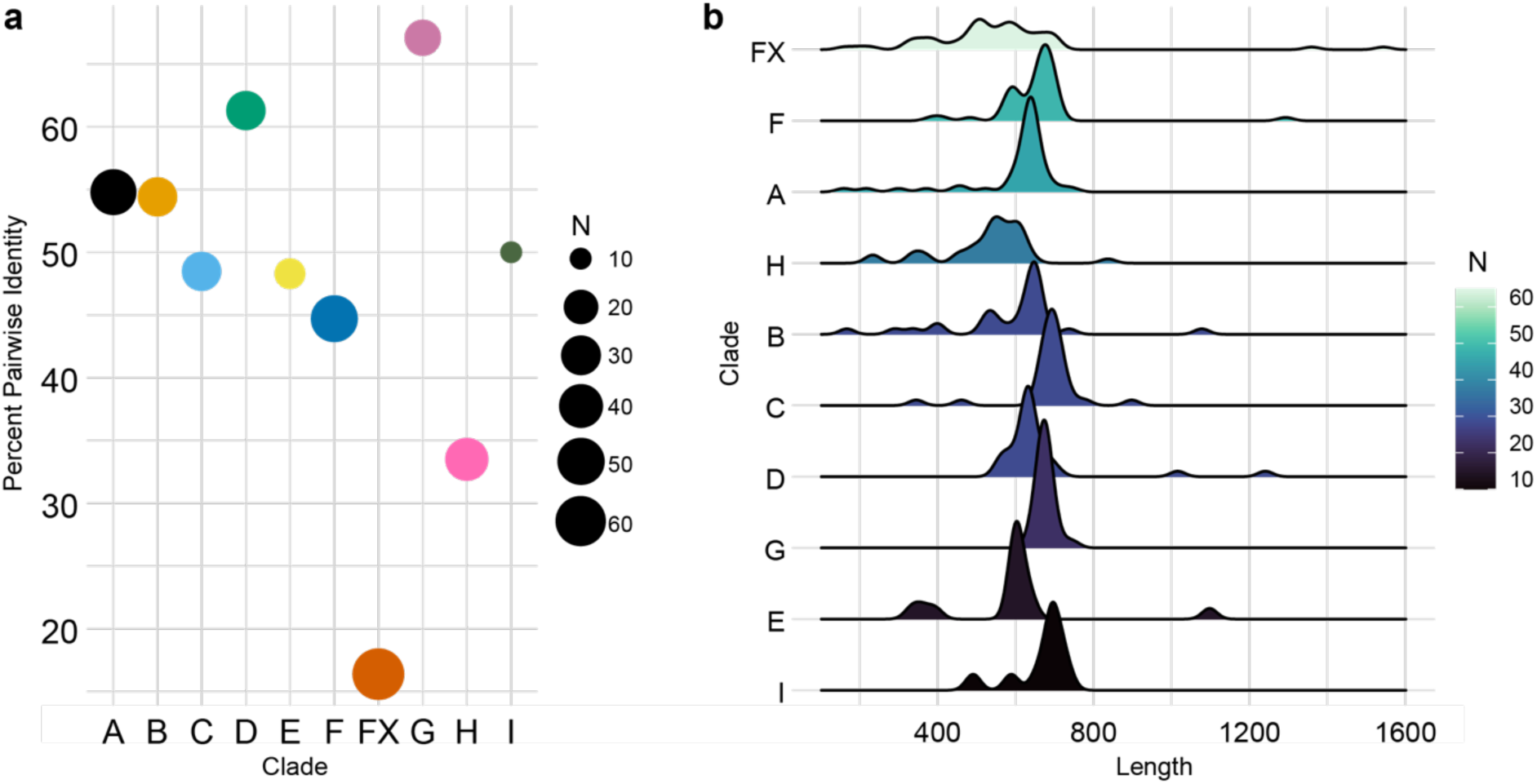
The EXO70FX clade is the most divergent and has the greatest variance in protein length compared to any other clade. All EXO70 proteins from fourteen monocot species shown in Figure 2 were compared for intra-clade sequence diversity and length. a) Percent pairwise identity for each clade based on intra-clade structure-based alignments. b) Protein lengths of members for each clade shown with density plots.

**Fig. S2.**
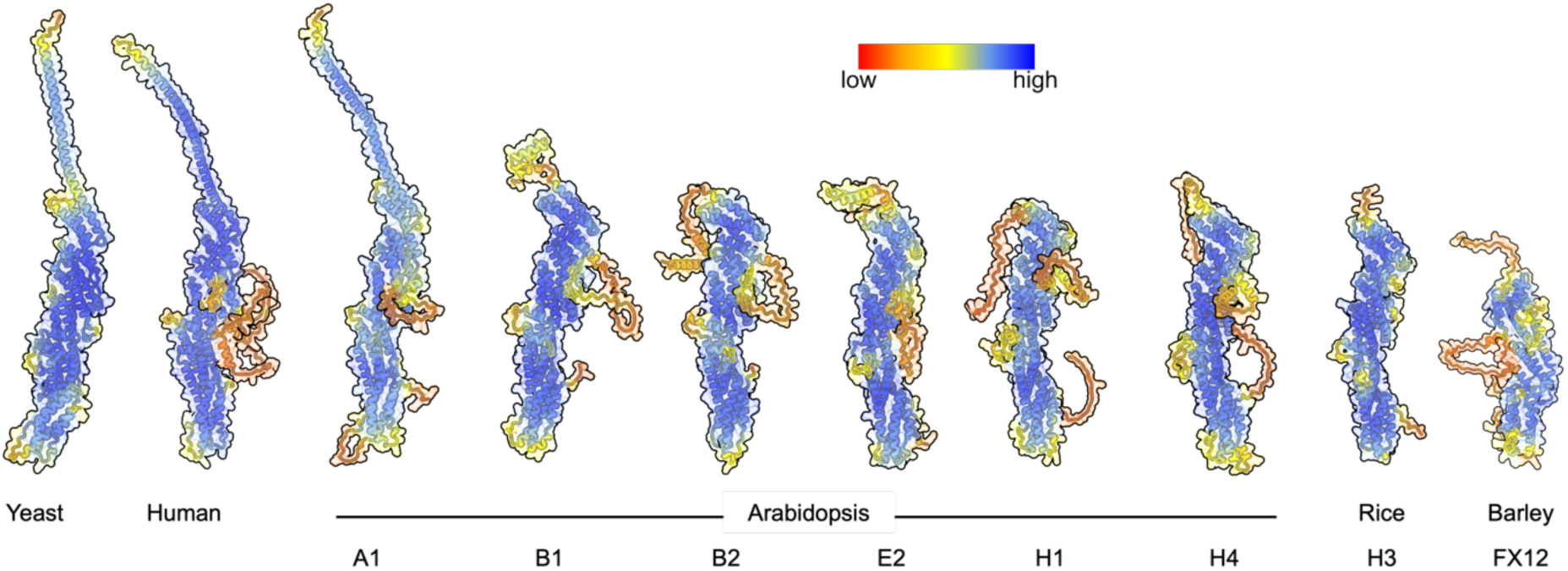
Exocyst-interacting EXO70s have high-confidence structural predictions. EXO70 structures of yeast and human, and selected plant EXO70s were predicted with AlphaFold2. Colouring by pLDDT indicates general high per-residue confidence score in all models in predicted alpha helices. Disordered N-terminal regions and side chains are predicted with less confidence.

**Fig S3.**
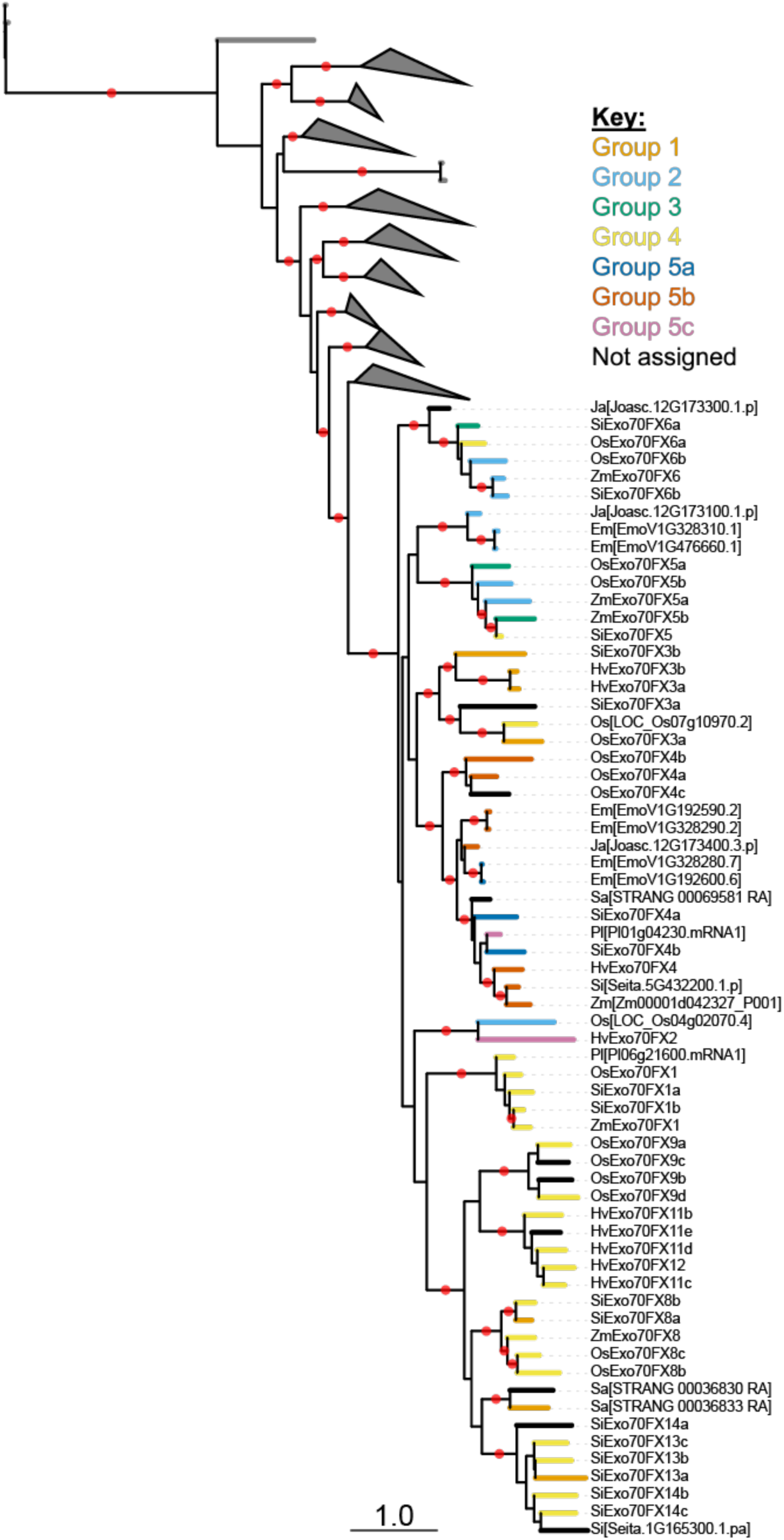
EXO70FX structural groups are distributed across Graminids. Reduced structure-based phylogeny from Figure 2 highlighting the 65 EXO70FX proteins. Branches are labelled by structural category. Red dots indicate bootstrap support greater than or equal to 80%. Scale indicates 1.0 substitution per site.

**Fig. S4.**
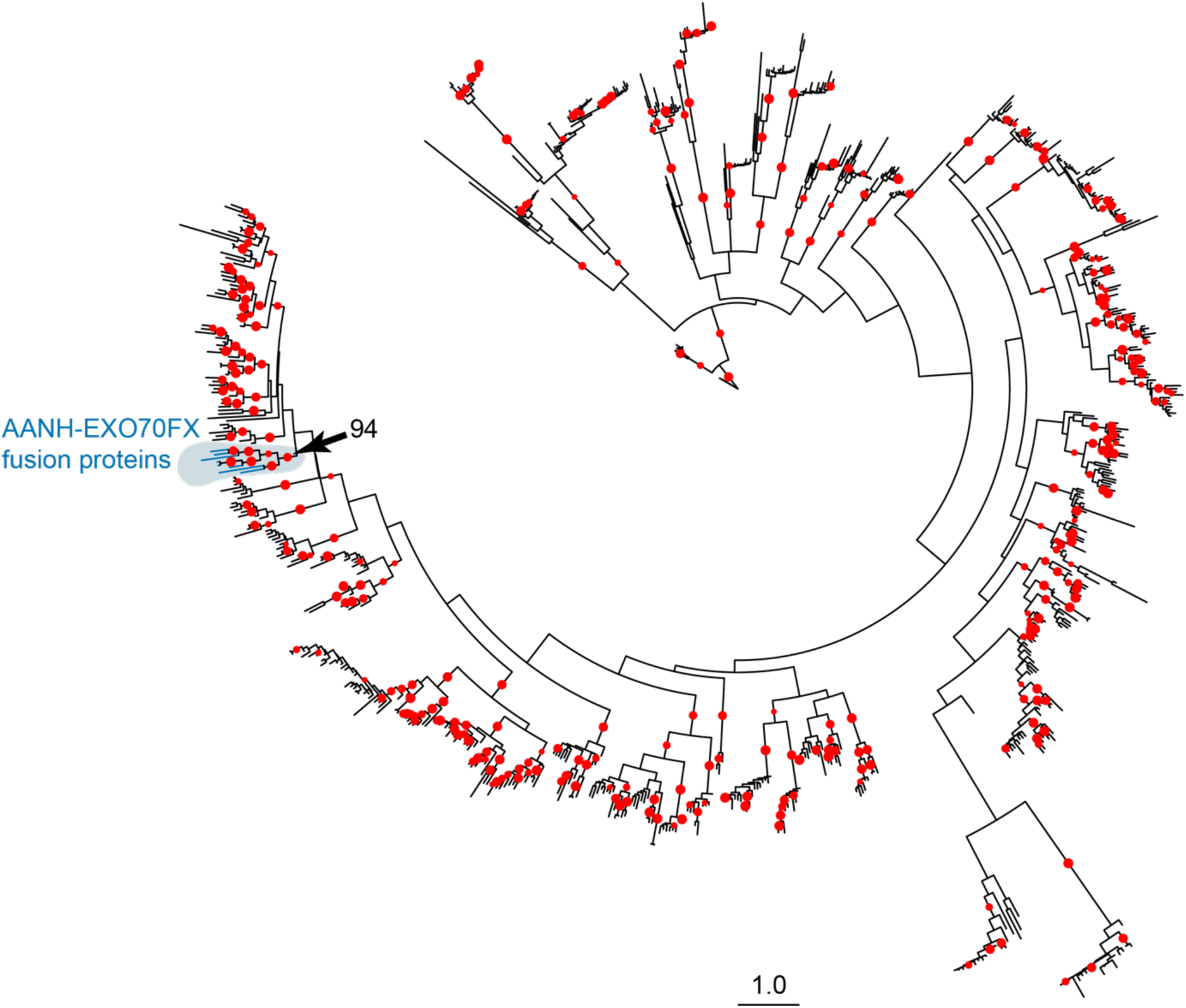
Adenine nucleotide alpha hydrolases-like (AANH)-EXO70FX fusion proteins are derived from a common ancestor. Structure-based maximum likelihood phylogenetic tree of 876 AANH domains derived from deposited PDB structures and 14 Poales species: *Hordeum vulgare*, *Triticum aestivum*, *Brachypodium distachyon*, *Oryza sativa*, *Sorghum bicolor*, *Setaria italica*, *Oropetium thomaeum*, *Zea mays*, *Ecdeiocolea monostachya*, *Joinvillea ascendens*, *Carex cristatella*, *Carex scoparia*, *Juncus effusus*, and *Juncus inflexus*. Branches of each of the eight AANH-EXO70FX proteins identified in Fig. 3b are shown in blue and highlighted in grey. The EXO70FX-fused AANH domains are derived from class VI plant U-box proteins (PUBs), which are a family of U-box-protein kinase fusion proteins (Trenner et al. 2022). Bootstrap support of greater than or equal to 80% is shown at branch midpoints with a red dot. The monophyletic AANH-EXO70FX clade indicated with an arrow is supported at 94%. Scale indicates 1.0 substitution per site.

**Fig S5.**
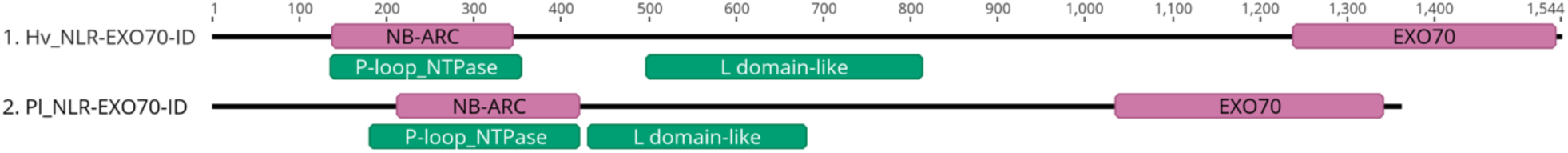
Two EXO70FX members are integrated domains in NLRs. Barley and stalkgrass (*Pharus latifolius*) NLR proteins with integrated EXO70 domains, HORVU.MOREX.r3.2HG0098620.1 and Pl01g04230.mRNA1, respectively. As identified by InterProScan, PFAM domains are shown in pink, and relevant SUPERFAMILY domains are shown in green.

**Fig. S6.**
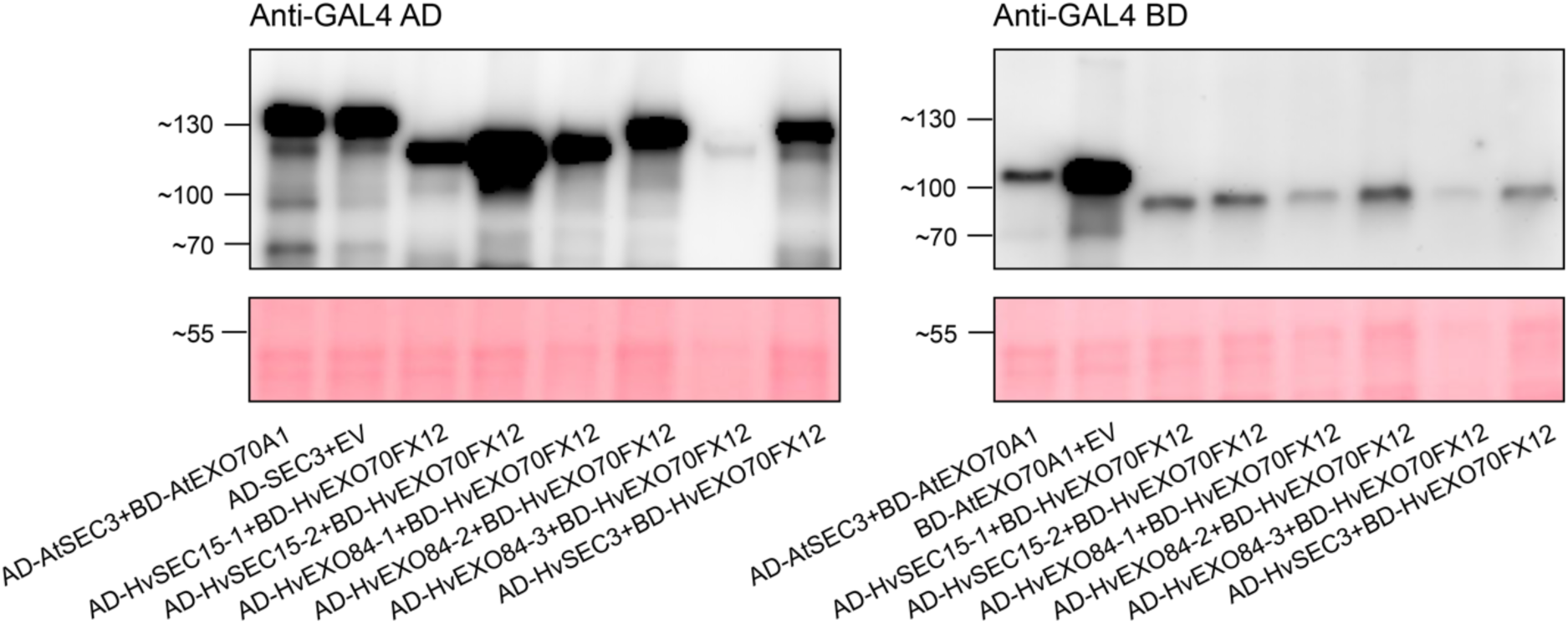
All exocyst subunits accumulate in Y2H. When fused to the GAL4 Activation Domain and expressed in yeast, the following proteins accumulated: AtSEC3 (121 kDA), HvSEC15-1 (109kDa), HvSEC15-2 (110 kDa), HvEXO84-1 (106 kDa), HvEXO84-2 (108 kDa), HvEXO84-3 (106 kDa), and HvSEC3 (122 kDa). When fused to the GAL4 DNA Binding Domain, AtEXO70A1 (91 kDa) and HvEXO70FX12 (76 kDa) accumulated. Three replicates were performed with similar results.

**Fig. S7.**
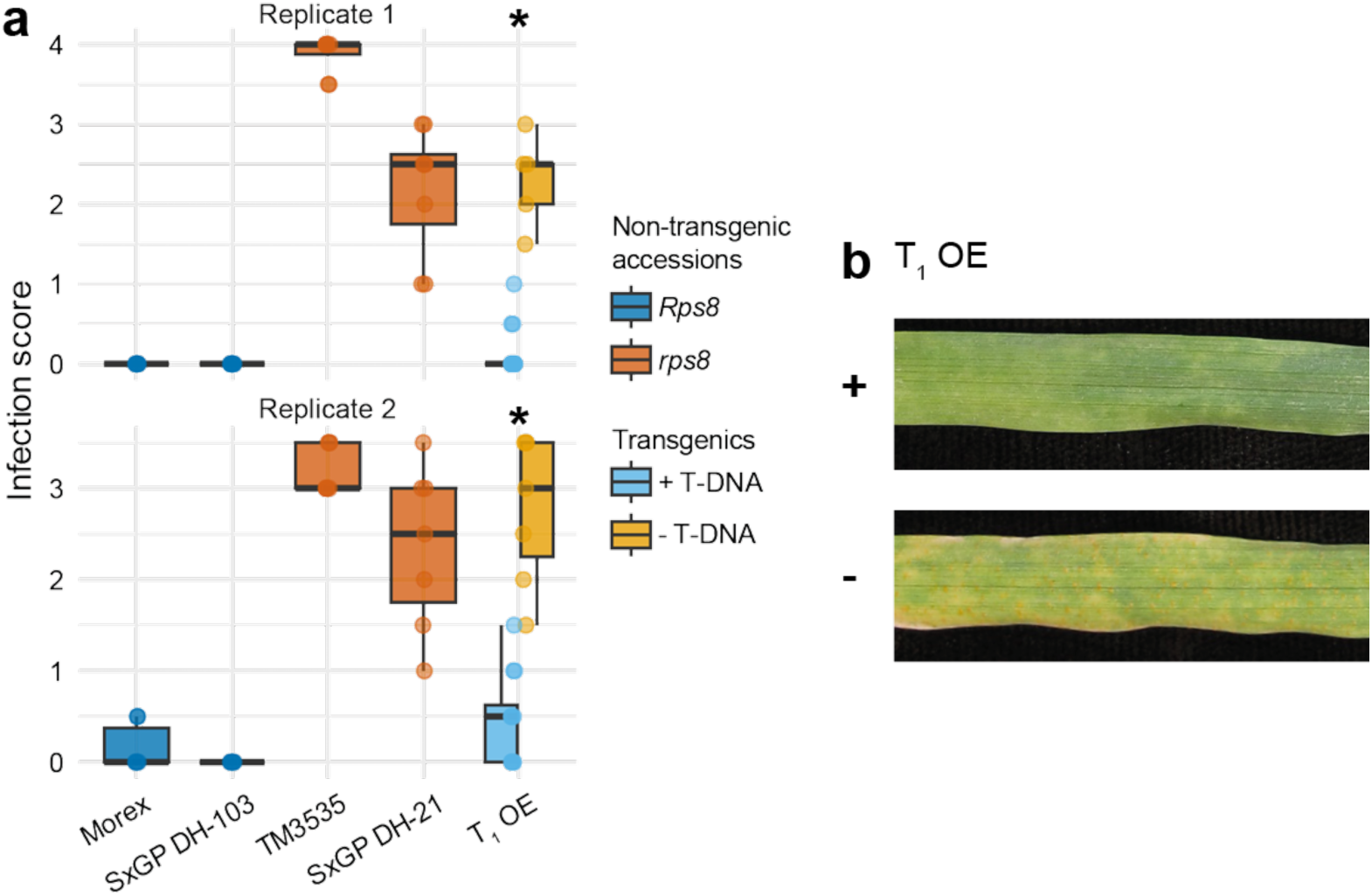
N-terminal tagging of HvEXO70FX12 does not abrogate function in transgenic barley. a) *Puccinia striiformis* f. sp. *tritici* isolate 20/092 infection scores are plotted for non-transgenic controls and a segregating T_1_ family derived from a hemizygous parent co-expressing *HvPur1* and *3*×*Flag-HvExo70FX12*. *Rps8* indicates the presence of a functional locus encoding *HvPur1* and *HvExo70FX12*, while *rps8* accessions lack resistance. Progeny of the transgenic family express resistance that significantly co-segregates with presence of T-DNA in two replicates based on the Wilcoxon-Mann-Whitney (WMW) test (replicate 1: W = 80, p-value = 0.00019; replicate 2: W = 83.5, p-value = 0.00043). In each replicate, approximately 16 progeny were tested. b) Photographs show an example line carrying T-DNA (+) expressing resistance, whereas a line without the T-DNA (-) is susceptible. Photographs were taken of leaves 15 days after inoculation.

**Fig. S8.**
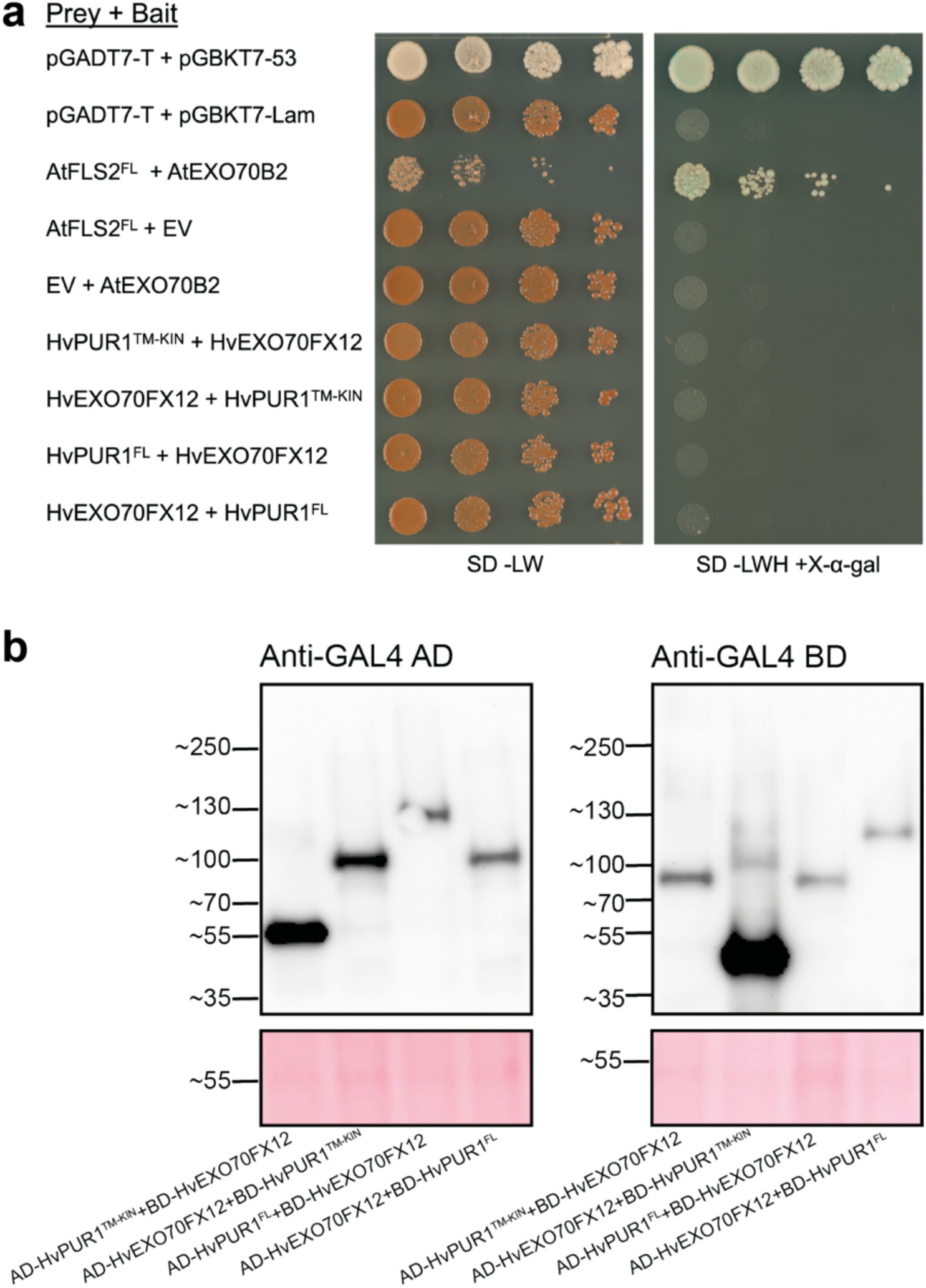
HvEXO70FX12 and HvPUR1 do not interact in Y2H. a) Matchmaker® Gold Y2H positive (pGADT7-T + pGBKT7-53) and negative (pGADT7-T + pGBKT7-Lam) controls were used in addition to AtFLS2-AtEXO70B2, which have been previously demonstrated to interact (Wang et al., 2020). Growth on synthetic defined (SD) -Leu/-Trp media indicates presence of activation and binding domain plasmids in yeast, while growth on SD/-Leu/-Trp/- His/+X-α-Gal media indicates interaction of bait and prey proteins. Three replicates were performed with similar results. b) HvEXO70FX12 (∼75 kDa), full-length HvPUR1 (∼130 kDa), and the HvPUR1 truncated transmembrane-intracellular region (TM-KIN; ∼55 kDa) accumulate in yeast in both bait and prey constructs. Two replicates were performed with similar results.

**Fig. S9.**
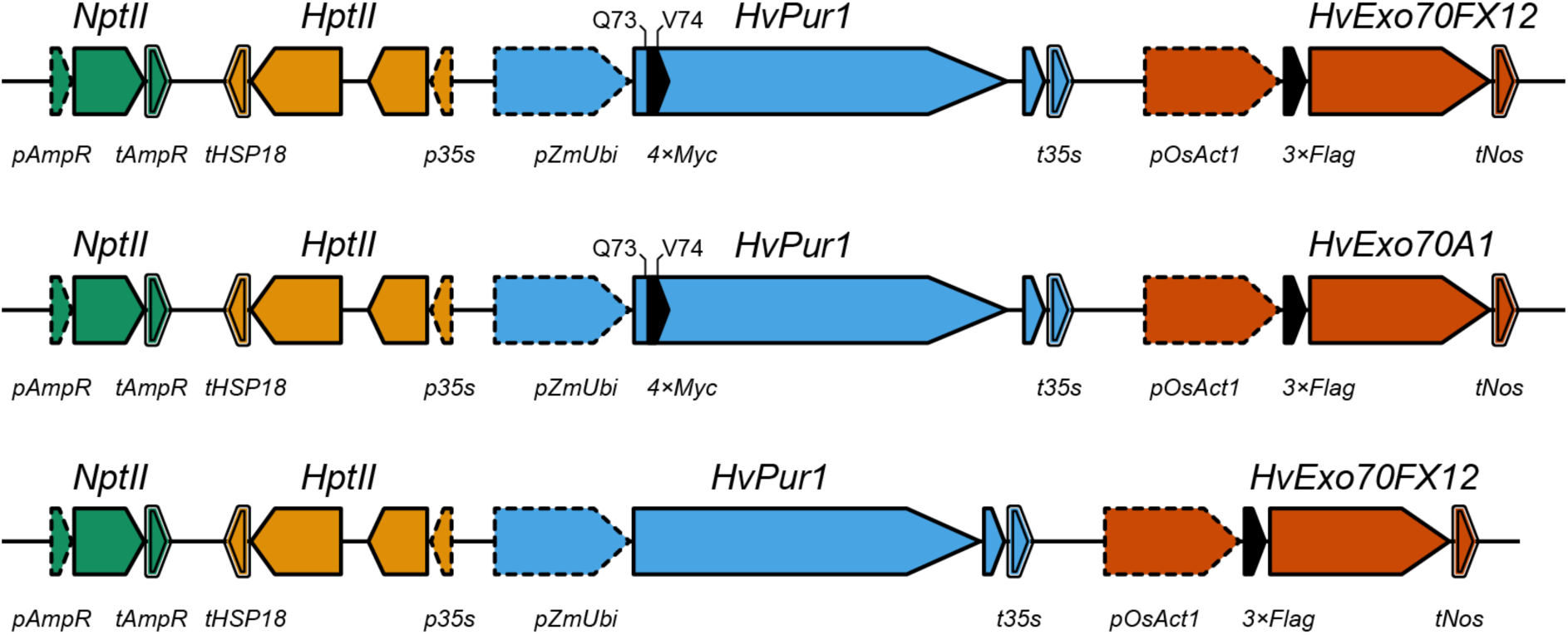
Gene constructs used to generate transgenic barley for proteomics and HvEXO70FX12 N-terminal tag functional complementation. The barley accession SxGP DH-47, which lacks the presence of *Pur1* and *Exo70FX12*, was transformed with three-gene transgenic cassettes. The schematic depicts the promoters (dashed outline), terminators (double outline), coding sequences (solid outline), and tags (black fill) for each transgenic cassette. Cassettes for proteomics included selection marker hygromycin under control of the CaMV+TMV *35s* promoter and *AtHSP18* terminator; *HvPur1* with an embedded *4xMyc* tag between HvPUR1^73Q^ and HvPUR1^74V^ under the control of *ZmUbi* promoter and CaMV *35s* terminator; and either *HvExo70FX12* or *HvExo70A1* fused to an N-terminal *3xFlag* tag under the control of the *OsAct1* promoter and *A. tumefaciens Nos* terminator. The native intron of *HvPur1* was maintained. The cassette for HvEXO70FX12 N-terminal complementation was created identically (bottom), except *HvPur1* was untagged. In each transgenic cassette, the hygromycin selection marker was in reverse orientation. The acceptor plasmid (pICSL4723) backbone contains kanamycin resistance conferred by *NptII*. Schematics are not to scale.

**Supplementary Table 1.** EXO70FX members predicted with AlphaFold2 and categorized based on domain structure.

| ID | Species | Structural Category |
| --- | --- | --- |
| HORVU.MOREX.r3.7HG0711570.1 | *Hordeum vulgare* | 1 |
| HORVU.MOREX.r3.7HG0711580.1 | *Hordeum vulgare* | 1 |
| LOC_Os07g10920.1 | *Oryza sativa* | 1 |
| Seita.1G130000.1.p | *Setaria italica* | 1 |
| Seita.4G068100.1.p | *Setaria italica* | 1 |
| Seita.5G035600.1.p | *Setaria italica* | 1 |
| STRANG_00036833-RA | *Streptochaeta angustifolia* | 1 |
| EmoV1G328310.1 | *Ecdeiocolea monostachya* | 2 |
| EmoV1G476660.1 | *Ecdeiocolea monostachya* | 2 |
| Joasc.12G173100.1.p | *Joinvillea ascendens* | 2 |
| LOC_Os01g28600.1 | *Oryza sativa* | 2 |
| LOC_Os01g67810.1 | *Oryza sativa* | 2 |
| LOC_Os04g02070.4 | *Oryza sativa* | 2 |
| Seita.5G419600.1.p | *Setaria italica* | 2 |
| Zm00001d040755_P001 | *Zea mays* | 2 |
| Zm00001d042461_P001 | *Zea mays* | 2 |
| LOC_Os02g36619.1 | *Oryza sativa* | 3 |
| Seita.5G419400.1.p | *Setaria italica* | 3 |
| Zm00001d031682_P001 | *Zea mays* | 3 |
| HORVU.MOREX.r3.2HG0208920.1 | *Hordeum vulgare* | 4 |
| HORVU.MOREX.r3.2HG0208940.1 | *Hordeum vulgare* | 4 |
| HORVU.MOREX.r3.2HG0208970.1 | *Hordeum vulgare* | 4 |
| HORVU.MOREX.r3.4HG0407730.1 | *Hordeum vulgare* | 4 |
| LOC_Os01g67820.1 | *Oryza sativa* | 4 |
| LOC_Os05g30640.1 | *Oryza sativa* | 4 |
| LOC_Os05g30660.1 | *Oryza sativa* | 4 |
| LOC_Os06g08460.1 | *Oryza sativa* | 4 |
| LOC_Os07g10970.2 | *Oryza sativa* | 4 |
| LOC_Os08g13570.1 | *Oryza sativa* | 4 |
| LOC_Os09g17810.1 | *Oryza sativa* | 4 |
| PI06g21600.mRNA1 | *Pharus latifolius* | 4 |
| Seita.4G050300.1.p | *Setaria italica* | 4 |
| Seita.4G050500.1.p | *Setaria italica* | 4 |
| Seita.4G068000.1.p | *Setaria italica* | 4 |
| Seita.4G232500.1.p | *Setaria italica* | 4 |
| Seita.4G285400.1.p | *Setaria italica* | 4 |
| Seita.4G285500.1.p | *Setaria italica* | 4 |
| Seita.4G285800.1.p | *Setaria italica* | 4 |
| Seita.5G175300.1.p | *Setaria italica* | 4 |
| Zm00001d015616_P001 | *Zea mays* | 4 |
| Zm00001d050462_P001 | *Zea mays* | 4 |
| EmoV1G192600.6 | *Ecdeiocolea monostachya* | 5a |
| EmoV1G328280.7 | *Ecdeiocolea monostachya* | 5a |
| Seita.5G432000.1.p | *Setaria italica* | 5a |
| Seita.5G432100.1.p | *Setaria italica* | 5a |
| EmoV1G192590.2 | *Ecdeiocolea monostachya* | 5b |
| EmoV1G328290.2 | *Ecdeiocolea monostachya* | 5b |
| HORVU.MOREX.r3.3HG0310710.1 | *Hordeum vulgare* | 5b |
| Joasc.12G173400.3.p | *Joinvillea ascendens* | 5b |
| LOC_Os07g10910.1 | *Oryza sativa* | 5b |
| LOC_Os07g10940.1 | *Oryza sativa* | 5b |
| Seita.5G432200.1.p | *Setaria italica* | 5b |
| Zm00001d042327_P001 | *Zea mays* | 5b |
| HORVU.MOREX.r3.2HG0098620.1 | *Hordeum vulgare* | 5c |
| PI01g04230.mRNA1 | *Pharus latifolius* | 5c |

**Supplementary Table 2.**
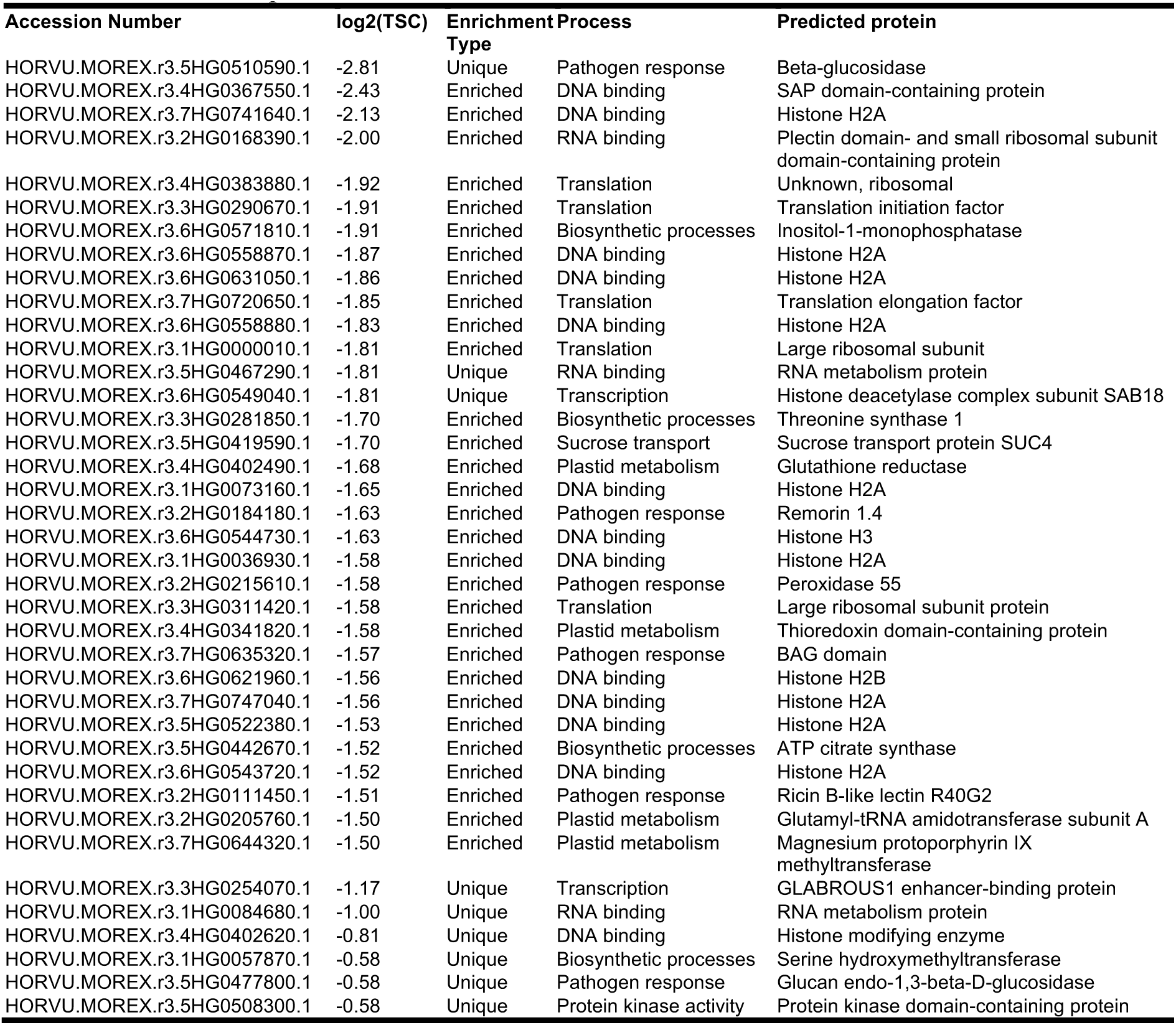
All unique or enriched (≥2.8X) proteins associated with HvEXO70FX12 compared to HvEXO70A1.

